# Receptor binding may directly activate the fusion machinery in coronavirus spike glycoproteins

**DOI:** 10.1101/2021.05.10.443496

**Authors:** Yuzhang Wang, Lucy Fallon, Lauren Raguette, Stephanie Budhan, Kellon Belfon, Darya Stepanenko, Stephanie Helbock, Sarah Varghese, Carlos Simmerling

**Affiliations:** Laufer Center for Physical and Quantitative Biology, Stony Brook University; Department of Chemistry, Stony Brook University, Stony Brook University; Department of Applied Mathematics and Statistics, Stony Brook University; Department of Materials Science and Chemical Engineering, Stony Brook University; Undergraduate Program in Biology, Stony Brook University

## Abstract

SARS-CoV-2, the causative agent of the COVID-19 pandemic, is an enveloped RNA virus. Trimeric spike glycoproteins extend outward from the virion; these class I viral membrane fusion proteins mediate entry of the virus into a host cell and are the dominant antigen for immune response. Cryo-EM studies have generated a large number of structures for the spike either alone, or bound to the cognate receptor ACE2 or antibodies, with the three receptor binding domains (RBDs) seen closed, open, or in various combinations. Binding to ACE2 requires an open RBD, and is believed to trigger the series of dramatic conformational changes in the spike that lead to the shedding of the S1 subunit and transition of the spring-loaded S2 subunit to the experimentally observed post-fusion structure. The steps following ACE2 binding are poorly understood despite extensive characterization of the spike through X-ray, cryo-EM, and computation. Here, we use all-atom simulations, guided by analysis of 81 existing experimental structures, to develop a model for the structural and energetic coupling that connects receptor binding to activation of the membrane fusion machinery.

## Introduction

A coronavirus spike is a homotrimeric glycoprotein, with each protomer consisting of two subunits S1 and S2; both are heavily decorated with glycans.(1) The N-terminal S1 subunits sit atop the spike and are responsible for recognizing and binding a host cell receptor and stabilizing the S2 core.(2-9) In SARS-CoV-2, each S1 subunit consists of an N-terminal domain (NTD), a receptor binding domain (RBD), and two C-terminal domains (CTD1 and CTD2); the S1/S2 interface lies at the C-terminal end of CTD2 (**Figure S1**).(10-12) The RBD in the S1 subunit is responsible for recognizing and binding angiotensin converting enzyme 2 (ACE2) (**Figure 1**).(3, 6-9, 13) The RBD alternates between at least two main conformational states relative to the remainder of the spike: ‘open’ and ‘closed’.(11, 12, 14, 15) An open RBD is a prerequisite for ACE2 binding; in the closed state binding of ACE2 is precluded by a steric clash with the RBDs of other protomers.(12, 14, 16, 17)

**Figure 1:**
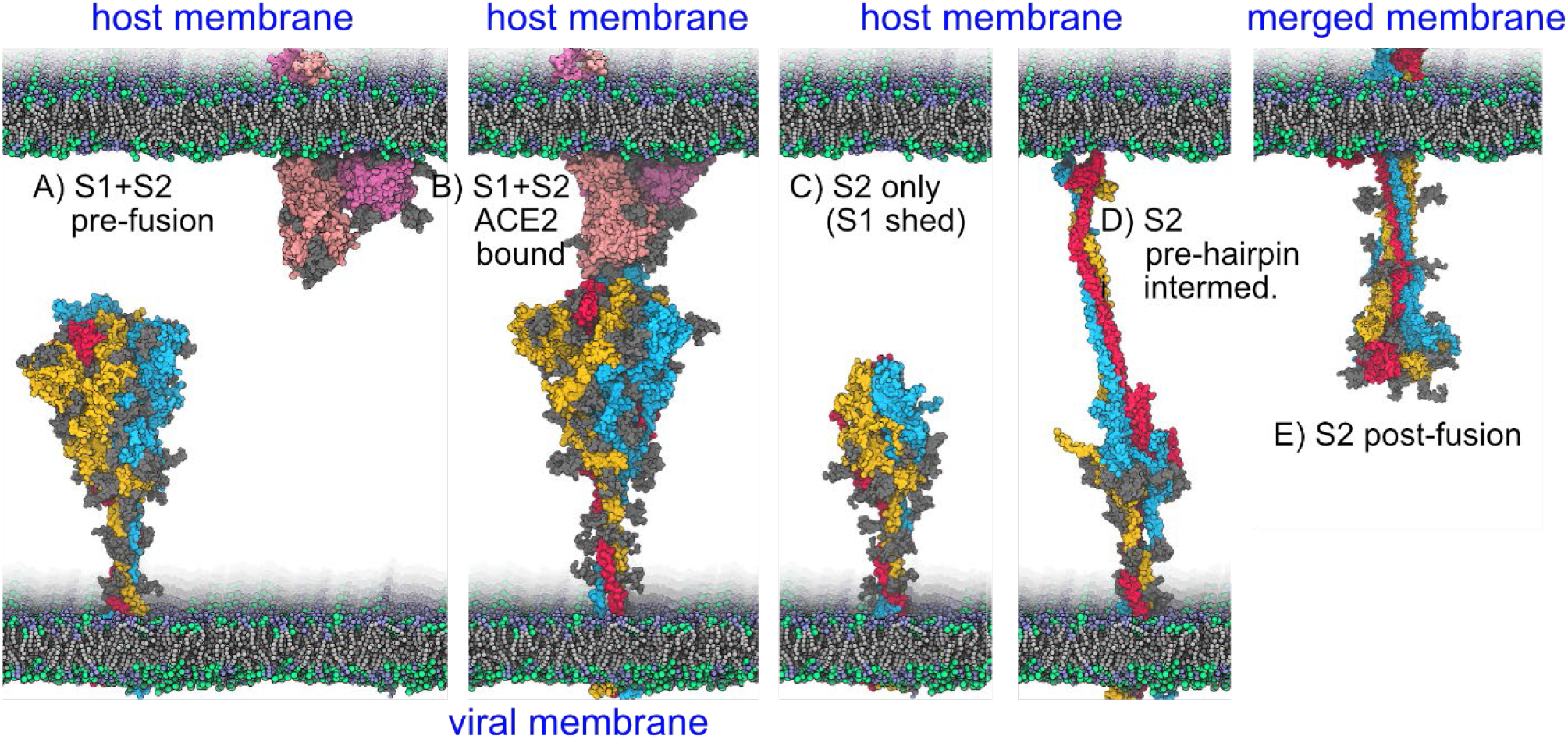
Structure models along the presumed pathway for spike-mediated membrane fusion in SARS-CoV-2. Each spike protomer is shown in a different primary color, ACE2 is shown in pink, and glycans are indicated in gray. A) The pre-fusion spike contains both S1 and S2 subunits. B) The spike binds to ACE2 using an open RBD. C) The S1 subunit sheds, leaving a metastable S2. D) The HR1 are released, extending the CH coiled-coil and inserting the FP into the host cell membrane. E) HR1 and HR2 come together to form a 6-helix bundle, merging the viral and host membranes. *Only states A, B and E have been observed experimentally; C and D are models to suggest general features*.

The fusion machinery of the spike is contained in the S2 subunit.(18) The S2 is composed of an upstream helix (UH) at the N-terminus, the fusion peptide proximal region (FPPR), the fusion peptide (FP), the first conserved heptad repeat region (HR1), a long central α-helix (CH), and a connector domain (CD) that leads into the C-terminal ‘stalk’ which contains the second heptad repeat (HR2), and the transmembrane domain (TM) that spans the viral membrane, followed finally by an intracellular region (**Figure S1**).(11, 12, 19)

The SARS-CoV-2 spike possesses two distinct cleavage sites that must both be processed by host proteases to foster efficient membrane fusion (S1/S2 and S2’).(4, 5, 20-22). The priming site at the S1/S2 boundary disconnects the S1 and S2 subunits, permitting eventual S1 dissociation (shedding). Cleavage at the S1/S2 site in SARS-CoV-2 is thought to occur during the secretory pathway; this is possible due to insertion of a dibasic furin recognition sequence (_682_RRAR_685_) that is not present in SARS and many other coronaviruses(21). The activation site S2’ (_814_KRSF_817_)(20, 23) is in the S2 domain at the FPPR/FP boundary, and is believed to be cleaved by membrane-bound host proteases such as TMPRSS2(23-26) or cathepsin(23, 27).

At some point after the S1 region binds ACE2 (**Figure 1B**), the S1 subunits dissociate to expose the metastable S2 core (**Figure 1C**), which is “spring-loaded” similar to influenza hemagluttinin(28). S2 subsequently undergoes large-scale refolding that initiates membrane fusion.(18, 19, 29-31) S2’ cleavage releases the fusion peptides at the end of HR1, although the timing of cleavage relative to ACE2 binding and S1 shedding remains unclear. The FP diffuse to and insert into the host cell membrane, presumably forming a short-lived pre-hairpin intermediate tethering the two membranes (**Figure 1D**).(18, 29, 32) This structure refolds again to adopt the post-fusion structure, with a six-helix bundle(23, 29, 33) formed from the HR1 and HR2 helices that are attached to the spike stalk and fusion peptides, respectively (**Figure 1E**). These dramatic rearrangements allow one or more spikes to bring the two lipid bilayers into contact, generating a fusion pore and releasing the virion’s contents into the host cell.(29)

What couples the receptor-binding function of the S1 subunit to the membrane fusion machinery of the S2 subunit? The “ratcheting” mechanism(34) suggests that opening of multiple RBDs leads to significant loss of S1-S2 contact area, leading to S1 shedding, after which the metastable S2 trimer refolds(29). However, experiments have imaged the pre-fusion spike bound to 1, 2 or 3 ACE2 proteins **(Figure 2**), suggesting that RBD ratcheting is insufficient to induce S1 shedding, even with S1/S2 cleavage(6). It is plausible that cleavage at S2’ is a prerequisite for S1 shedding, but the S2’ recognition motif appears inaccessible to proteases in pre-fusion spike structures. Other host factors may participate in spike-mediated viral entry(35-37), but in vitro S1 shedding and adoption of post-fusion structure can occur in their absence(19, 29).

**Figure 2.**
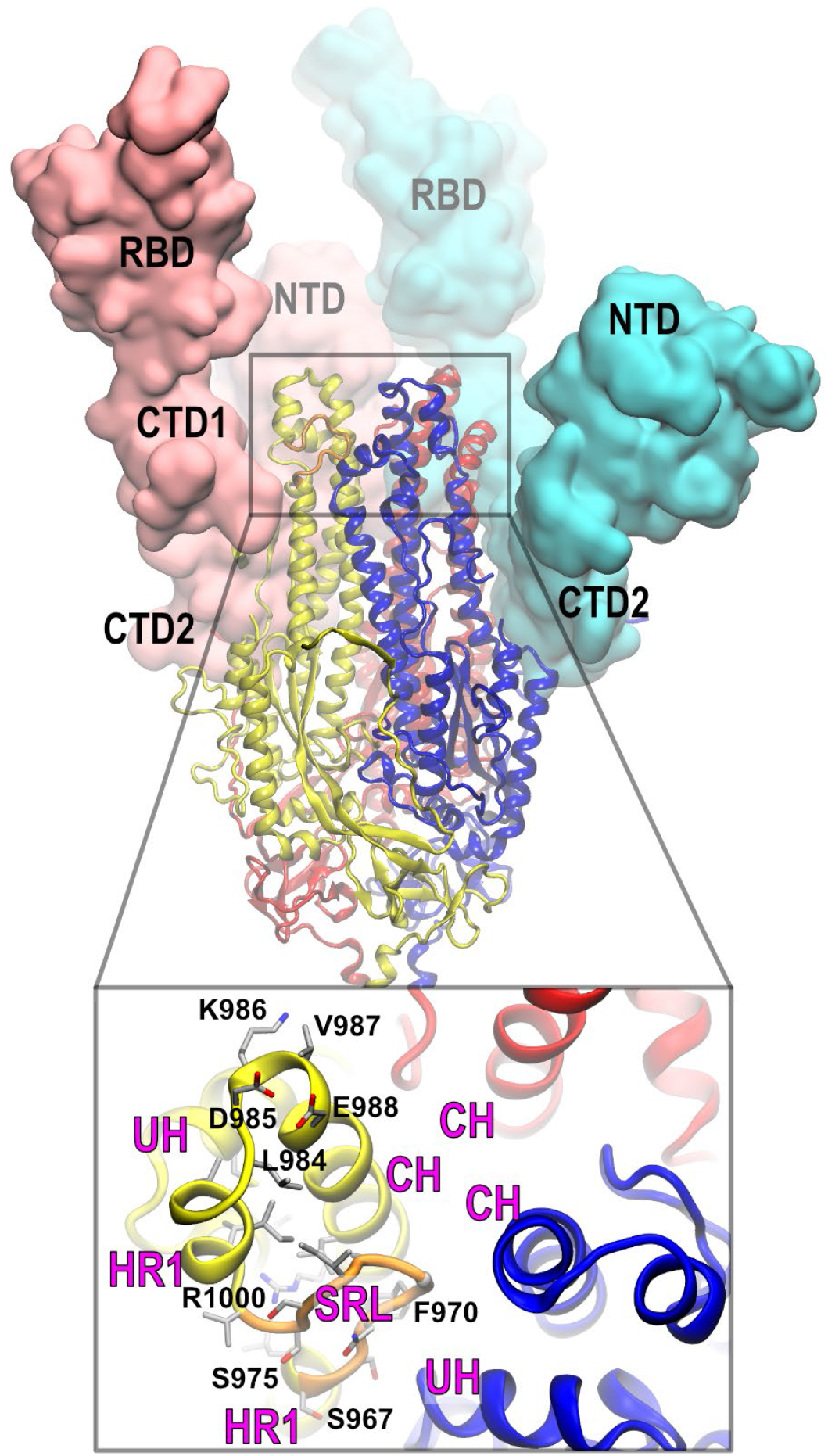
Structure of the pre-fusion spike ectodomain with 3 open RBDs (7CAK(38)). S1 subunits are shown in a space-filling model and S2 are drawn with ribbons. For clarity the S1 subunit is not shown for one protomer (yellow). Each protomer is a different primary color, with one S2 loop highlighted in orange. Inset: close-up view of the top of the S2 subunit for one protomer in wild-type 6XR8.

One possible explanation for these observations is that all available experimental structures with more than one open RBD employed sequences with stabilizing amino acid substitutions at the N-terminal of the CH α-helix (such as the widely employed “2P” spike(39), K986P/V987P). The enhanced stability of the 2P spike has led to many valuable studies of spike structure and function, as well as COVID-19 vaccines. The 2P modification is intended to block CH extension, since proline is relatively rigid and lacks the ability to donate backbone hydrogen bonds needed to extend the CH α-helix.(39)

In this work we describe a model for how receptor binding may trigger significant conformational changes in the S2 fusion machinery prior to S1 shedding. In our model, lengthening of the central coiled-coil occurs *prior to* S1 shedding; this shares features with a reported intermediate state in influenza hemagglutinin, in which α-helix extension in HA2 occurs prior to separation from HA1.(6) An important role is played by an unusual loop in the S2 subunit, which we call the serine-rich loop (SRL) since four of ten amino acids are serine (S_**967**_SNFGAISSV_**976**_). The SRL is located between the 3^rd^ and 4^th^ α-helices of HR1, near the location of the 2P substitution at the top of the CH (**Figures 2, S2**). In addition to providing a hypothesis for events immediately following spike-receptor binding, the model also suggests caution when interpreting data obtained with stabilizing substitutions in the context of the spike-mediated fusion mechanism.

## Results

### 1: Analysis of experimental structures of the SARS-CoV-2 spike from the PDB

We selected representative experimental structures in four different states: 6XR8, 6VSB, 7CAK, and 6XRA. Although most structures include the 2P stabilizing substitutions(39), a few studies did not (referred to here as “wild-type”). These include the 6XR8(19) pre-fusion structure with all three RBDs closed, and the 6XRA(19) post-fusion structure. We also include the 6VSB(11) 2P-spike with 1 RBD open (“1-up”), and the 7CAK(38) 2P-spike with all 3 RBDs open (“3-up”) and bound to antibody Fab proteins (7CAK has high similarity to the 3-up 7A98(6) structure with each RBD bound to an ACE2 domain). Due to limited resolution in cryo-EM models, analysis and visualization of these four structures is supplemented with data extracted from 81 SARS-CoV-2 spike structures obtained from the PDB (80 pre-fusion and 1 post-fusion, data in **Figure S6**, with PDB codes in **Table S1**). Histograms of distances between the RBD and the top of CH or HR1 are shown in **Figure S6a**. In all structures, R815 at the S2’ cleavage site appears inaccessible to proteases and closely packed against A871, with α-carbon distances of 5.9 ± 0.3 Å in the 80 pre-fusion spike structures (**Figure S6b**).

**Figure 2** shows the 3-up spike, where the open RBDs expose the top of the S2 subunit trimer. This region contains the three central helices (CH) at the core, with each protomer forming a helix-turn-helix motif (HTH) connecting HR1 to CH. The top of the wild-type CH exhibits features of an N-capping motif(40), with the D985 side chain hydrogen bonding to the backbone N of E988 and hydrophobic contacts stabilizing a turn that leads from HR1 into the CH (**Figure 2**). The HTH are surrounded by the three upstream helices (UH); the upper horizontal α-helix of each UH forms a close contact with the CH of the same protomer.

#### An unusual loop is present near the top of S2

Moving from the HTH motif down the spike along HR1, two of the four α-helical segments of the pre-fusion HR1 (helix-3 and helix-4) are interrupted by the serine-rich loop. The SRL adopts a “C” shape, ending in nearly the same location as it started (**Figure 2**), with the α-carbon atoms of S967 and S975 separated by 5.4 ± 0.2 Å in the PDB structure set (**Figure S6c**).

The pre-fusion SRL has several unusual features. The segment from N969 to A972 forms a type-I’ β-turn (turn type *ad*(41)), where amino acids in the *a* and *d* positions both adopt positive φ backbone dihedral values. G971 occupies the *d* position, but the *a* position is occupied by the conserved F970, which likely is strained due to this disallowed backbone conformation (adopted in all pre-fusion structures but not in the post-fusion state, **Figure S6d**). The side chain of F970 is packed tightly against G999 in the CH (**Figure 2**, with a distance of 3.8 ± 0.2 Å, **Figure S6e**). Glycine is uncommon inside helices due to the additional entropic penalty of adopting a defined conformation, but any side chain at this position would clash with the F970 ring.

Another presumably strained feature of the pre-fusion SRL is the placement of R1000 inside a cluster of hydrophobic side chains capped by the SRL (**Figures 2, S2**). Arginine has a pKa near 14, and almost certainly remains charged even when buried in a nonpolar environment(42-44). The buried guanidino group is able to form only two hydrogen bonds (one to each side of the pocket), with backbone O atoms of I742 on the UH and S975 on the SRL (PDB data in **Figures S6f, g**). Cation burial may serve as a reservoir to store spike folding energy for release during the transition to the solvent-exposed environment observed in the post-fusion spike.

In the closed spike, each HR1-CH turn is covered by the RBDs of both other protomers (**Figure S3**), with the counterclockwise protomer contacting the CH and clockwise protomer contacting HR1. Specific interactions are shown in **Figure S4**. D428 is in the proximity of CH K986 (P986 in the 2P spike), potentially forming a favorable interaction that stabilizes the closed RBD(19). The RBD of the counterclockwise protomer forms extensive interactions with the top of HR1 and the turn connecting it to CH, including a hydrogen bond between the S383 sidechain OG and the backbone of D985, and the backbone N of S383 with the backbone O of R983 (PDB data in **Figure S6a**). The side chain amine of K386 on nearly every closed RBD forms a C-capping interaction with HR1, stabilizing the exposed backbone O of I981 (**Figure S6h**). Overall, these contacts with the two RBDs help stabilize the ends of both α-helices in the HTH.

#### RBD opening results in extensive loss of contacts between the S1 and S2 subunits

Our representative systems for the open spike include the 1-up 6VSB, and 3-up 7CAK (**Figure 2**); both include the stabilizing 2P substitution. As expected, the contacts between the RBD and the S2 HTH motif are lost when the RBD opens (**Figure 2**). However, additional S1-S2 interactions lower down the spike are also disrupted. As the RBD opens, the attached CTD1 domain moves outward(6), though to a lesser extent than the RBD. This shift results in loss of contacts between side chains on the CTD1 and the short upper helix of HR1, including R567-D979 and D571-S975. The outward shift of CTD1 leaves this HR1 segment completely free of contact with the S1 subunit, even more so when the RBD is engaged by ACE2 or antibodies (**Figure S5**). Data extracted from the 80 pre-fusion structures support a strong correlation between RBD opening and loss of CTD1-HR1 contacts (**Figure S6i**,**j**).

#### Comparing the pre-fusion and post-fusion spike reveals important changes in the region near the SRL

In the post-fusion state, the S1 subunit has dissociated and only the S2 subunit is present (**Figure 1E**). The most obvious conformational change is the dramatic extension of CH, in which the HTH motif, SRL and the remainder of HR1 rotate upward to form a single continuous helix with CH, which adopt the coiled-coil motif that extends the FP toward the host cell membrane.(19, 29) The process may be driven in part by energetically favorable changes; compared to the pre-fusion spike, R1000 becomes solvent-exposed, F970 adopts an allowed backbone conformation (**Figure S6d**), hydrophobic side chains from the HR1/SRL form a tightly packed inter-protomer hydrophobic cluster, and six new inter-protomer salt bridges are formed within a stretch of three turns of α-helix (**Figure S7**).

Three of these salt bridge pairs (D994 - R995’) are close to the site of the SRL in the pre-fusion structure. The SRL is deeply inserted between the UH and CH of neighboring protomers, and the SRL appears to sterically block R995 from approaching D994 (**Figure S8**; with a broad distribution of D994 - R995’ distances in PDB structures, **Figure S6k**). Relocation of the SRL in the post-fusion structure is associated with closer approach of the UH and CH, permitting formation of the D994-R995’ salt bridges (**Figure S8**). The other three new inter-protomer salt bridge pairs in the post-fusion structure (R983-D985’) involve amino acids from the upper short helix-4 of HR1 which are distant in the pre-fusion structure but become neighbors in the coiled-coil of the post-fusion state (**Figure S7**).

#### The upper S2 rotates relative to the lower S2 during membrane fusion

(45) Despite the dramatic differences between the pre-fusion and post-fusion structures, the S2 structure near the base of the CH shows minimal changes. Overlap on this region highlights a significant twist in S2 that accompanies the transition, with the upper CH and UH rotating about a fulcrum that is highly localized near M731 and E1017 (**Figure S9**). The net result is that the post-fusion S2 becomes smaller and more tightly packed than in the pre-fusion spike; these changes may be blocked until the SRL wedge is removed from between CH and UH (**Figure 3**). The distance between N751 α-carbon pairs is reduced from 31.4 ± 0.5 Å in the PDB pre-fusion structures **(Figure S6l**) to 26 Å in the post-fusion spike.

**Figure 3.**
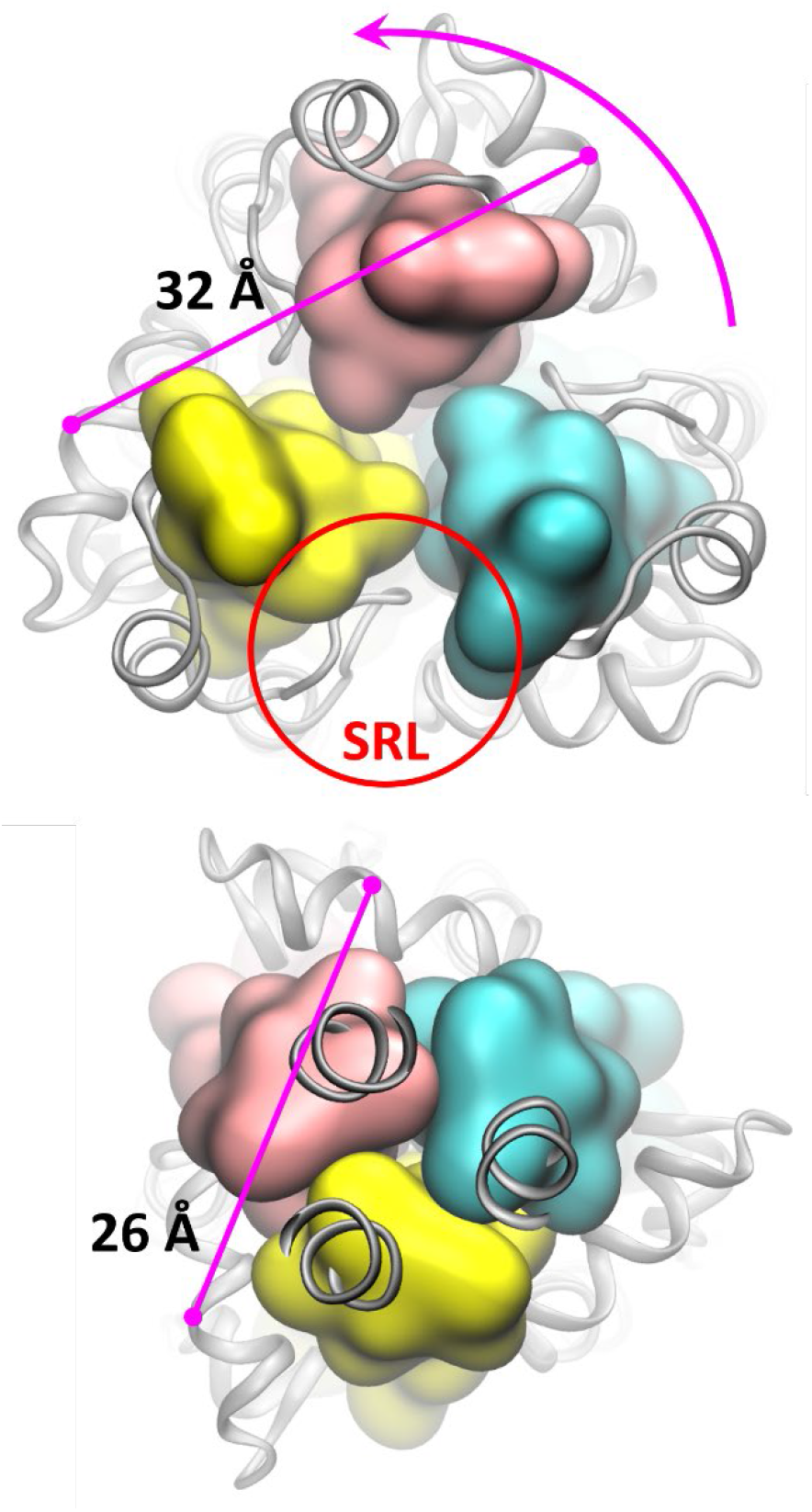
Comparison of the S2 core in the pre-fusion (upper, 6XR8) and post-fusion (lower, 6XRA) spike. Amino acids on the pre-fusion CH (V987 to A1015) are shown as space-filling. The distance between N751 on neighboring UH for one pair of protomers is indicated with a pink line. After removal of the SRL, rotation and repacking of S2 results in a notable size reduction in the S2 subunit.

#### A proposed role for the SRL in triggering of spike-mediated membrane fusion

Unless the SRL serves a functional purpose, one would expect these strained amino acids (e.g. F970, G999, R1000) to be replaced during viral evolution. What mechanistic role could necessitate their conservation? When the RBDs open, the packing around the upper helix-4 of HR1 is released, while the HR1 helix-3 below the SRL remains constrained by the S1 CTD1 (**Figure S5**).

We propose that RBD opening may provide the space needed to rotate upward the short HR1 helix-4, extending the CH by 2 turns. The N-terminal of the lengthened helix is able to remain connected to the more-distant HR1 helix-3 through unfolding of the compact SRL. SRL unfolding could be compensated energetically by relaxation of F970, solvent exposure of R1000, and formation of the D994-R995’ salt bridges, while adding the HR1 helix-4 to the CH may be driven by formation of the hydrophobic cluster and R983-D985’ salt bridges (**Figure S7**). The released energy may compensate for the disruption of interactions with S1 as the S2 core rotates and becomes more compact (**Figure 3**). *Thus, the SRL-mediated lengthening of the CH coiled-coil may precede, and even facilitate, S1 shedding by releasing energy via relaxation of multiple strained structural elements*.

Why are these changes not seen in the current open-RBD experimental structures? Each HTH motif connecting CH and HR1 is covered by two different RBDs (**Figure S3**). This suggests that a single open RBD is insufficient to trigger changes in CH; requiring multiple open RBDs could have a functional importance by preventing spurious spike activation when a single RBD opens to search for a target receptor. Only when a single RBD is held open for significant periods (such as by ACE2 binding) does it become likely for a second RBD to open(6, 34). Experimental structures such as 7CAK display multiple open, bound RBDs, however it is plausible that stabilization of the top of the CH via the 2P substitution also suppresses the motion of the upper HR1. Fewer structures have been solved for the wild-type pre-fusion spike (16 of 80 in our data set); all are either closed or 1-up. One study observed the 2-up unbound wild-type spike on inactivated virions, but upon purification only the closed form was present and a wild-type 2-up structure was not refined.(15)

Even with open RBDs, S2 triggering may be slow due to the need to disrupt the HTH hydrophobic cluster and CH N-cap, leading to a kinetically controlled process as in hemagglutinin(46). pH also has been suggested to play a role in RBD opening(47). pH may also be a factor in the kinetics of CH extension; protonation of Asp weakens N-capping and can give rise to pH-dependent protein conformational changes(48). The D985 N-cap may be particularly sensitive due to an expected upward shift in pKa from the close proximity of E988 (**Figure 2**). Overall, the role of pH in spike-mediated membrane fusion remains to be elucidated.

Is it reasonable for these changes in S2 to take place *prior* to S1 shedding? A confounding factor in the analyses presented above is that the experimental pre-fusion and post-fusion structures have many other differences, primarily the presence or absence of the entire S1 subunit. It remains unclear if a partially extended coiled-coil would fit under open RBDs. If so, is the length of the SRL sufficient to retain the link to helix-3 in HR1 as the coiled-coil becomes more distant? Where could the unfolded SRL be accommodated? Furthermore, would the coiled-coil and new salt bridges be able to form within the constraints imposed by the S1 shell? Since the currently available experimental structures do not address these questions, we supplement existing knowledge using model building and all-atom simulations.

### 2. Simulation results

#### Standard MD simulations of the closed and 3-up spike ectodomain

Following protocols in our previous work(49), we prepared models of the wild-type spike glycoprotein in explicit water in 2 conformations: with all three RBDs either closed (based on 6XR8) or open (based on 7CAK). Details are provided as Supplementary Information. We carried out four independent, unrestrained simulations of ∼ 400 ns for each conformation. Both systems were reasonably stable, and no significant changes in the top of S2 or the SRL were observed (RMSD values of 0.5 – 1.5 Å compared to the 6XR8 closed spike, **Figure S10**).

In all simulations, a water molecule diffused from the bulk solvent into the hydrophobic pocket under each SRL, bridging the R1000 NE and the backbone O of S968; water density analysis confirmed localization of water at this position in all three protomers (**Figure S11**). No change in local geometry was needed, suggesting that this site may be occupied by a water molecule in cryo-EM experiments. This is consistent with reports that arginine in nonpolar environments often retains contact to water microdroplets; the affinity of the first water molecule to guanidinium has been estimated to be 10-11 kcal/mol.(50)

#### Model systems probe longer timescale dynamics of S2

Following the pioneering work of Carr and Kim in which peptide fragments were used to reveal the spring-loaded mechanism of hemagglutinin(28), we created two smaller model systems using different subsets of the pre-fusion S2 subunit (shown in **Figure S12**). In the “medium” system, we retained the central components of the trimer, including the CH, SRL, HR1 and UH (M731-K776, D950-R1019). All amino acids below the fulcrum of S2 rotation **(Figure S9)** were removed, and neutral termini were added and restrained during MD. In the “small” model, we further truncated the medium system by deleting the SRL and lower HR1, from D950 to S975, with neutral termini added but no restraints applied to V976. Both systems were simulated in full atomic detail with explicit water.

We began with simulations of the small model in order to determine if a single α-helix is the preferred conformation of the HTH motif. During simulations of 1.5 μsec in explicit water for the wild-type sequence, the HR1 helix-4 in all three protomers spontaneously rotated upwards to form a continuous helix with CH. The final structure closely matches the conformation of the same region in the post-fusion spike 6XRA, with RMSD values decreasing from the initial ∼ 18 Å to ∼ 2 Å (**Figure S13**). These results suggest that the pre-fusion helix-turn-helix motif at the top of the S2 subunit is locally strained, and in the absence of the SRL, the system spontaneously relaxes the HTH to a structure in which CH and HR1 form a continuous α-helix. This shows strong similarity to the spring-loaded mechanism(51) in hemagglutinin.

Importantly, control simulations of the same model system using the 2P substitution showed no extension of the CH in any of the protomers, with RMSD values remaining near 15 Å during 4 μsec MD (**Figure S13**). This dramatically contrasts with the results obtained with the wild-type model, supporting our hypothesis that the 2P spike may be hindered from sampling the HR1 rotation.

We next asked if the SRL, once unfolded, could serve as a linker from HR1 helix-3 to the top of the extended CH. Initial MD simulations of the medium model were run for 400 ns, in which the pre-fusion structure was stable and no upward rotation of the short HR1 helix was observed. The difference from the small model MD suggests that unfolding the SRL, or perhaps breaking the hydrogen bond between R1000 and S975 (discussed above) may contribute a significant kinetic barrier; therefore, we used steered molecular dynamics (SMD) to achieve CH extension (See Methods). Over the course of a 40ns simulation, the α-carbon atoms of each helix-turn-helix (using N978 - E1017) were steered to match the structure for this region obtained from MD on the small model. The SRL (S967-L977) was allowed to move freely, and the remainder of the system was weakly restrained to the initial post-fusion structure. After SMD, an additional 1 μsec of MD was carried out without any restraints except at the truncated base; the extended helices were stable during this simulation. The final structure is shown in **Figure S14**.

In the pre-fusion structure, the CH is N-capped by D985, with hydrophobic side chains on HR1 packing against CH (**Figure 2**). After extension of CH in the medium model, a highly similar N-capping motif formed, including hydrogen bonding from the side chain of N978 to the exposed backbone NH of I980/L981 at the new helix N-terminal, and the hydrophobic V976 and/or L977 stabilizing the turn leading down the SRL to HR1 (**Figure S15**). The lower HR1 helix remains connected to the top of the extended CH via unfolding of the SRL (**Figure S15**). The placement of the unfolded SRL alongside the CH in our model is comparable to an extended segment of HR2 that follows the same path in the post-fusion structure, with F970 in a similar location as F1156 in the post-fusion spike (**Figure S16**).

#### Simulations of CH extension using the full spike ectodomain

The S2-only model systems cannot provide insight into whether the observed process of SRL unfolding and CH extension can take place *prior* to S1 shedding, where it must occur inside the steric constraints of the 50 Å-wide(6) cylinder formed by the NTDs and open RBDs of S1 (**Figure 2**). No significant changes to S2 were seen in the 3-up standard MD simulations described above, so we steered the 3-up pre-fusion spike to adopt the final conformation of the SRL and HTH motif that were obtained from the medium model system. Next, we refined a pathway for CH extension in the full spike by using the nudged elastic band method (NEB), with a protocol similar to what we employed(49) to map the RBD opening pathway. Here, the endpoints were defined as the 3-up spike before and after CH extension. We did not aim to find a globally optimal path revealing the exact order of events for such a complex change, but simply to ensure that a reasonable transition pathway was possible without significant steric clash with S1. Snapshots during CH extension are shown in **Figure S17**; the short HR1 helix rotates upward as the SRL unfolds. The NEB pathway mapping was followed by a fully unrestrained, 250 ns MD simulation on the spike with extended CH. A video file is included as supplementary information.

As with the model systems, the RMSD of the HTH compared to the post-fusion spike was reduced from ∼ 18 Å to ∼ 3 Å as the SRL unfolded, where it remained during unrestrained MD (time traces shown in **Figure S18 A, E**). N978 forms an N-capping interaction with the CH in all three protomers, similar to the N-capping motif involving D985 in the pre-fusion structure. The hydrophobic cluster of I980/L984 side chains stabilizing the coiled-coil, and salt bridges between R983’/D985’ (**Figure 4**) also formed as the CH lengthened (**Figure S18B-D**). Prior to CH extension, the D994/R995’ pairs sampled a broad distance distribution similar to that in the pre-fusion PDB structures (**Figure S6L**), but closer distances corresponding to stable salt bridges are observed after unfolding of the SRL (**Figure S18B**).

**Figure 4.**
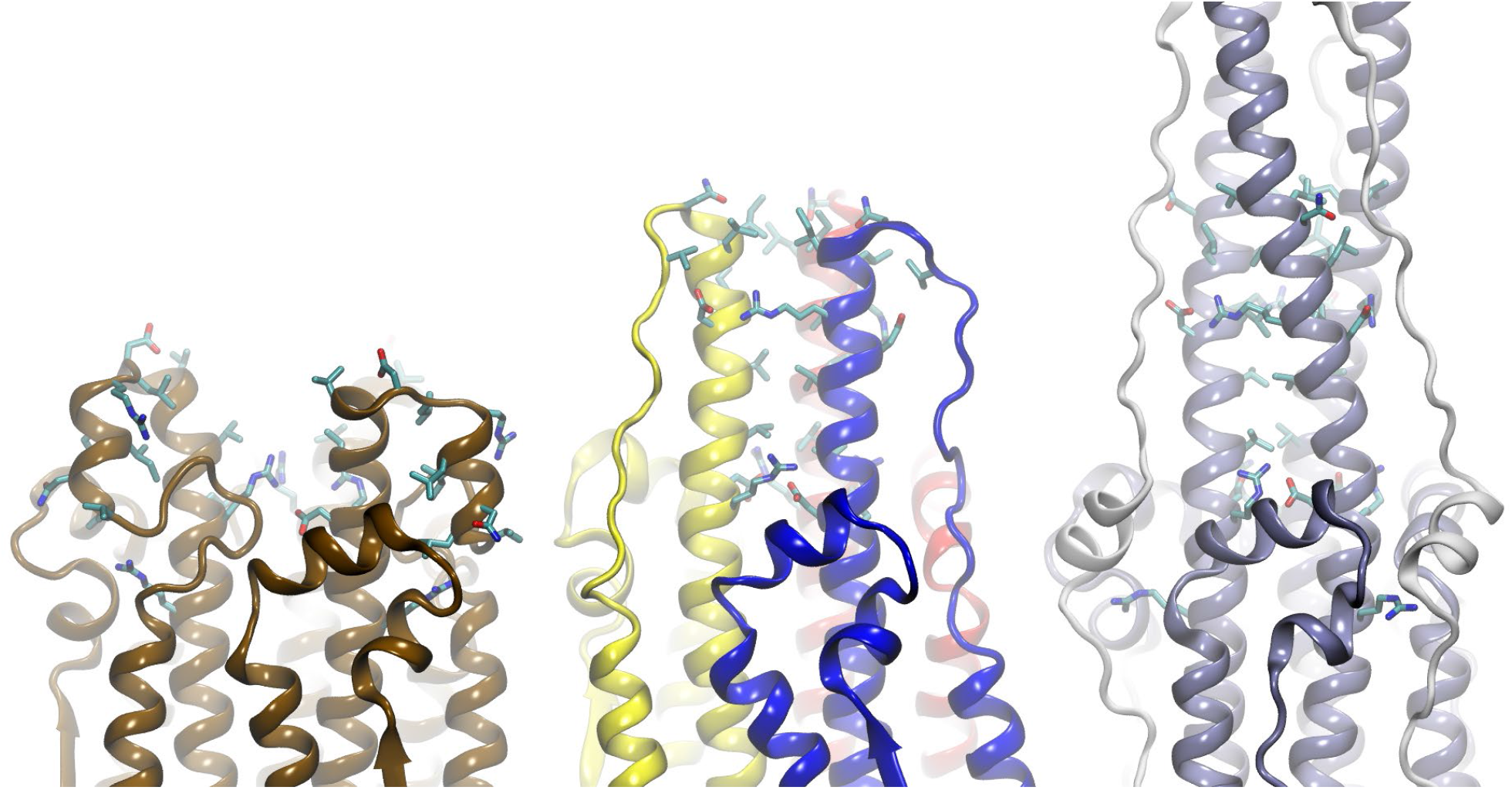
Comparison of features between the upper S2 subunit of the wild-type spike in the pre-fusion 6XR8 (left, copper), intermediate model (middle, primary colors), and post-fusion 6XRA (right, gray) structures. *The S1 subunits in the left and middle structures are present but omitted for clarity*; the post-fusion spike contains only S2. **Figure S15** shows these regions with labels on selected amino acids.

The upper portion of the S2 subunit in our intermediate state model is compared to the same region in the pre-fusion and post-fusion cryo-EM structures in **Figure 4**, and the intermediate structure including the S1 subunit with RBDs is shown in **Figure 5**. The comparison highlights the role of SRL unfolding in allowing the HR1 helix-4 to extend the CH, while maintaining a connection to the HR1 helix-3 that is held in place by S1.

**Figure 5:**
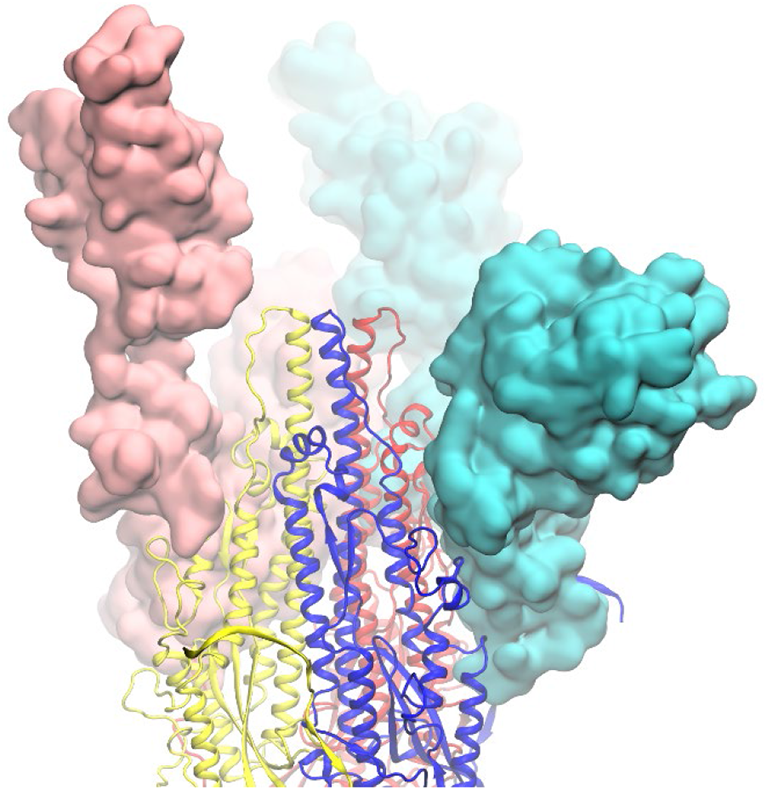
Structure of the 3-up spike with unfolded SRL and partially extended CH, comparable to the pre-fusion structure in **Figure 2**. Protomers are shown as primary colors, with the S1 subunits in space-filling and the S2 subunits as ribbons. As in **Figure 2**, the yellow S1 subunit is omitted to provide a clear view of the S2 subunit.

Importantly, R1000 is exposed to solvent upon SRL unfolding. We estimated the hydration free energy change of R1000 by applying a Poisson-Boltzmann (PB) approach to MD snapshots, with average per-protomer solvation free energies of -47 ± 1, -48 ± 1 and -58 ± 2 kcal/mol in the closed, 3-up, and 3-up extended-CH spike respectively **(Figure S19**). The data indicate that RBD opening does little to mitigate the poor environment of R1000, but significant energy is released upon SRL unfolding (∼ 10 kcal/mol per protomer). Exposing this buried cation to solvent may provide a significant energetic contribution to the “spring-loading” of the spike.

A notable difference in the pre-fusion and post-fusion spike is the rotation of the upper S2 relative to lower S2, presumably driven by relaxation in S2 packing once the SRL is removed from the wedge position between CH and UH (**Figures 3, S9**). We quantified the rotation using a dihedral angle for each UH; the angle is relatively stable in MD for the 3-up pre-fusion spike, but after SRL unfolding and extension of the CH, the UH slowly approach (but did not yet reach) the rotation adopted in the post-fusion spike (**Figure S18F**). This suggests that even the small change of extending the CH by two turns is sufficient to trigger more extensive rearrangements in S2 despite the constraints imposed by S1. A weakened S1-S2 interface may be readily disrupted by lateral forces on the S1 subunit when both spike and ACE2 are bound to their respective membranes(17), leading to S1 shedding. Future simulations on longer timescales may provide additional insight into the destabilization of the S1-S2 interface.

Our structural model suggests a possible experimental approach to probing the role of the SRL. F970 and G999 are favorably positioned to form a disulfide bond connecting the lower part of the SRL to the CH. We performed standard MD simulations of the F970C/G999C pre-fusion spike, and the disulfide bond was accommodated with minimal structure perturbation (**Figure S20**). In the context of ACE2 binding, a non-2P spike with the F970C/G999C disulfide may allow the observation of early changes in the HTH motif that are frustrated in the 2P spike, while still preventing the dissociation of the lower HR1 that is required for the complete conversion to the post-fusion structure.

## Conclusions

Based on data extracted from 81 experimental spike structures in the PDB, along with observations from all-atom simulations, we propose a mechanistic hypothesis for receptor-induced triggering of the S2 fusion machinery. A key component is the SRL, an unusual loop that is ubiquitous in coronavirus spike structures. Our model rationalizes the compact structure of the SRL, its placement between two otherwise continuous α-helical segments in HR1, and the roles of several nearby highly conserved amino acids. Upon receptor binding, unfolding of the SRL and upward rotation of the HR1 can extend the CH by an additional two turns of α-helix, with the extended SRL retaining the link between the longer helix and the HR1 that remains held in place by the CTD1 of the S1 subunit. This process may be driven by spike folding energy that is stored in highly strained structure elements near the SRL that relax following SRL unfolding. Our simulations suggest that commonly used stabilizing substitutions at the top of CH may hinder these conformational changes, providing a possible rationale for why they have not been observed in experimental spike structures.

Our analysis suggests a model for how the fusion core can be activated directly by binding of the spike to host receptors such as ACE2. However, many other aspects remain unclear, such as the role of pH, possible cooperativity between the protomers, the timing of S2’ site exposure and cleavage, and other important events in spike-mediated membrane fusion. A more detailed mechanism may facilitate the rational design of small molecules that could block these changes, or perhaps serve as a catalyst to trigger them prematurely, leading to irreversible activation of the spike and neutralization of the virus. The highly conserved fusion mechanism suggests that such approaches could have broad applicability to coronaviruses.

## Acknowledgments

The authors thank Rommie Amaro, Arvind Ramanathan, Ronit Freeman, Adrian Roitberg, Daniel Raleigh and Tom Kurtzman for helpful discussions. We are grateful to Bing Chen for early access to the 6XR8 spike structure and John Stone for help with an alpha release of VMD.

This work was funded in part by the SUNY Research Seed Grant Program, the Stony Brook University Office of the Vice President for Research, the Research Corporation for Science Advancement (COVID Initiative grant #27350), and NIH grant R01GM107104 (C.S.). This work used resources services, and support provided via the COVID-19 HPC Consortium (https://covid19-hpc-consortium.org/), which is a unique private-public effort to bring together government, industry, and academic leaders who are volunteering free compute time and resources in support of COVID-19 research. Additional computer time was provided by AiMOS at Rensselaer Polytechnic Institute.

## References

1. Y. Watanabe, J. D. Allen, D. Wrapp, J. S. McLellan, M. Crispin, Site-specific glycan analysis of the SARS-CoV-2 spike. Science 369, 330–333 (2020).

2. H. Zhang, J. M. Penninger, Y. Li, N. Zhong, A. S. Slutsky, Angiotensin-converting enzyme 2 (ACE2) as a SARS-CoV-2 receptor: molecular mechanisms and potential therapeutic target. Intensive Care Medicine 46, 586–590 (2020).

3. J. Shang et al., Structural basis of receptor recognition by SARS-CoV-2. Nature 581, 221–224 (2020).

4. M. Hoffmann et al., SARS-CoV-2 Cell Entry Depends on ACE2 and TMPRSS2 and Is Blocked by a Clinically Proven Protease Inhibitor. Cell 181, 271–280.e278 (2020).

5. J. Shang et al., Cell entry mechanisms of SARS-CoV-2. Proceedings of the National Academy of Sciences 117, 11727–11734 (2020).

6. D. J. Benton et al., Receptor binding and priming of the spike protein of SARS-CoV-2 for membrane fusion. Nature 588, 327–330 (2020).

7. Q. Wang et al., Structural and Functional Basis of SARS-CoV-2 Entry by Using Human ACE2. Cell 181, 894–904.e899 (2020).

8. R. Yan et al., Structural basis for the recognition of the SARS-CoV-2 by full-length human ACE2. Science (New York, N.Y.) 2, 1444–1448 (2020).

9. J. Lan et al., Structure of the SARS-CoV-2 spike receptor-binding domain bound to the ACE2 receptor. Nature 581, 215–220 (2020).

10. Y. Huang, C. Yang, X.-f. Xu, W. Xu, S.-w. Liu, Structural and functional properties of SARS-CoV-2 spike protein: potential antivirus drug development for COVID-19. Acta Pharmacologica Sinica 41, 1141–1149 (2020).

11. D. Wrapp et al., Cryo-EM structure of the 2019-nCoV spike in the prefusion conformation. Science (New York, N.Y.) 1263, 1260–1263 (2020).

12. A. C. Walls et al., Structure, Function, and Antigenicity of the SARS-CoV-2 Spike Glycoprotein. Cell 181, 281–292 (2020).

13. W. Tai et al., Characterization of the receptor-binding domain (RBD) of 2019 novel coronavirus: implication for development of RBD protein as a viral attachment inhibitor and vaccine. Cellular & Molecular Immunology 17, 613–620 (2020).

14. M. Gui et al., Cryo-electron microscopy structures of the SARS-CoV spike glycoprotein reveal a prerequisite conformational state for receptor binding. Cell Research 27, 119–129 (2017).

15. Z. Ke et al., Structures and distributions of SARS-CoV-2 spike proteins on intact virions. Nature 588, 498–502 (2020).

16. W. Song, M. Gui, X. Wang, Y. Xiang, Cryo-EM structure of the SARS coronavirus spike glycoprotein in complex with its host cell receptor ACE2. PLoS Pathog 14, e1007236 (2018).

17. E. P. Barros et al., The flexibility of ACE2 in the context of SARS-CoV-2 infection. Biophys J 120, 1072–1084 (2021).

18. F. Li, Structure, Function, and Evolution of Coronavirus Spike Proteins. Annual Rev Virol 3, 237–261 (2016).

19. Y. Cai et al., Distinct conformational states of SARS-CoV-2 spike protein. Science 369, 1586–1592 (2020).

20. S. Belouzard, V. C. Chu, G. R. Whittaker, Activation of the SARS coronavirus spike protein via sequential proteolytic cleavage at two distinct sites. Proceedings of the National Academy of Sciences of the United States of America 106, 5871–5876 (2009).

21. B. Coutard et al., The spike glycoprotein of the new coronavirus 2019-nCoV contains a furin-like cleavage site absent in CoV of the same clade. Antiviral Research 176, 104742–104742 (2020).

22. J. A. Jaimes, N. M. André, J. S. Chappie, J. K. Millet, G. R. Whittaker, Phylogenetic Analysis and Structural Modeling of SARS-CoV-2 Spike Protein Reveals an Evolutionary Distinct and Proteolytically Sensitive Activation Loop. J Mol Biol 432, 3309–3325 (2020).

23. T. Tang, M. Bidon, J. A. Jaimes, G. R. Whittaker, S. Daniel, Coronavirus membrane fusion mechanism offers a potential target for antiviral development. Antiviral Research 178, 104792 (2020).

24. M. C. Johnson et al., Optimized Pseudotyping Conditions for the SARS-COV-2 Spike Glycoprotein. Journal of Virology 94, e01062–01020 (2020).

25. L. M. Reinke et al., Different residues in the SARS-CoV spike protein determine cleavage and activation by the host cell protease TMPRSS2. PLOS ONE 12, e0179177 (2017).

26. M. Hoffmann, H. Kleine-Weber, S. Pöhlmann, A Multibasic Cleavage Site in the Spike Protein of SARS-CoV-2 Is Essential for Infection of Human Lung Cells. Molecular Cell 78, 779–784.e775 (2020).

27. J. K. Millet, G. R. Whittaker, Physiological and molecular triggers for SARS-CoV membrane fusion and entry into host cells. Virology 517, 3–8 (2018).

28. C. M. Carr, P. S. Kim, A spring-loaded mechanism for the conformational change of influenza hemagglutinin. Cell 73, 823–832 (1993).

29. A. C. Walls et al., Tectonic conformational changes of a coronavirus spike glycoprotein promote membrane fusion. Proc Natl Acad Sci U S A 114, 11157–11162 (2017).

30. C. Liu et al., Viral Architecture of SARS-CoV-2 with Post-Fusion Spike Revealed by Cryo-EM. bioRxiv 10.1101/2020.03.02.972927, 2020.2003.2002.972927 (2020).

31. C. Liu et al., The Architecture of Inactivated SARS-CoV-2 with Postfusion Spikes Revealed by Cryo-EM and Cryo-ET. Structure 28, 1218–1224.e1214 (2020).

32. I. G. Madu, S. L. Roth, S. Belouzard, G. R. Whittaker, Characterization of a Highly Conserved Domain within the Severe Acute Respiratory Syndrome Coronavirus Spike Protein S2 Domain with Characteristics of a Viral Fusion Peptide. Journal of Virology 83, 7411–7421 (2009).

33. S. Xia et al., Fusion mechanism of 2019-nCoV and fusion inhibitors targeting HR1 domain in spike protein. Cell Mol Immunol 10.1038/s41423-020-0374-2 (2020).

34. A. C. Walls et al., Unexpected Receptor Functional Mimicry Elucidates Activation of Coronavirus Fusion. Cell 176, 1026–1039 e1015 (2019).

35. T. M. Clausen et al., SARS-CoV-2 Infection Depends on Cellular Heparan Sulfate and ACE2. Cell 183, 1043–1057.e1015 (2020).

36. L. Cantuti-Castelvetri et al., Neuropilin-1 facilitates SARS-CoV-2 cell entry and infectivity. Science 370, 856 (2020).

37. J. L. Daly et al., Neuropilin-1 is a host factor for SARS-CoV-2 infection. Science 370, 861 (2020).

38. Z. Lv et al., Structural basis for neutralization of SARS-CoV-2 and SARS-CoV by a potent therapeutic antibody. Science 369, 1505 (2020).

39. J. Pallesen et al., Immunogenicity and structures of a rationally designed prefusion MERS-CoV spike antigen. Proceedings of the National Academy of Sciences 114, E7348–E7357 (2017).

40. L. Serrano, A. R. Fersht, Capping and α-helix stability. Nature 342, 296–299 (1989).

41. M. Shapovalov, S. Vucetic, R. L. Dunbrack, Jr., A new clustering and nomenclature for beta turns derived from high-resolution protein structures. PLOS Computational Biology 15, e1006844 (2019).

42. C. A. Fitch, G. Platzer, M. Okon, B. Garcia-Moreno E, L. P. McIntosh, Arginine: Its pKa value revisited. Protein Science 24, 752–761 (2015).

43. B. Roux, Lonely Arginine Seeks Friendly Environment. Journal of General Physiology 130, 233–236 (2007).

44. M. J. Harms, J. L. Schlessman, G. R. Sue, B. García-Moreno E, Arginine residues at internal positions in a protein are always charged. Proceedings of the National Academy of Sciences 108, 18954 (2011).

45. X. Fan, D. Cao, L. Kong, X. Zhang, Cryo-EM analysis of the post-fusion structure of the SARS-CoV spike glycoprotein. Nature Communications 11, 3618 (2020).

46. D. Baker, D. A. Agard, Influenza hemagglutinin: kinetic control of protein function. Structure 2, 907–910 (1994).

47. T. Zhou et al., Cryo-EM Structures of SARS-CoV-2 Spike without and with ACE2 Reveal a pH-Dependent Switch to Mediate Endosomal Positioning of Receptor-Binding Domains. Cell Host & Microbe 28, 867–879.e865 (2020).

48. Y. Huang et al., Helix N-Cap Residues Drive the Acid Unfolding That Is Essential in the Action of the Toxin Colicin A. Biochemistry 58, 4882–4892 (2019).

49. L. Fallon et al., Free Energy Landscapes for RBD Opening in SARS-CoV-2 Spike Glycoprotein Simulations Suggest Key Interactions and a Potentially Druggable Allosteric Pocket. ChemRxiv Preprint (2020).

50. B. Gao, T. Wyttenbach, M. T. Bowers, Protonated Arginine and Protonated Lysine: Hydration and Its Effect on the Stability of Salt-Bridge Structures. The Journal of Physical Chemistry B 113, 9995–10000 (2009).

51. C. M. Carr, C. Chaudhry, P. S. Kim, Influenza hemagglutinin is spring-loaded by a metastable native conformation. Proceedings of the National Academy of Sciences 94, 14306 (1997).

