## Supplemental Information for "Receptor binding may directly activate the fusion machinery in coronavirus spike glycoproteins"

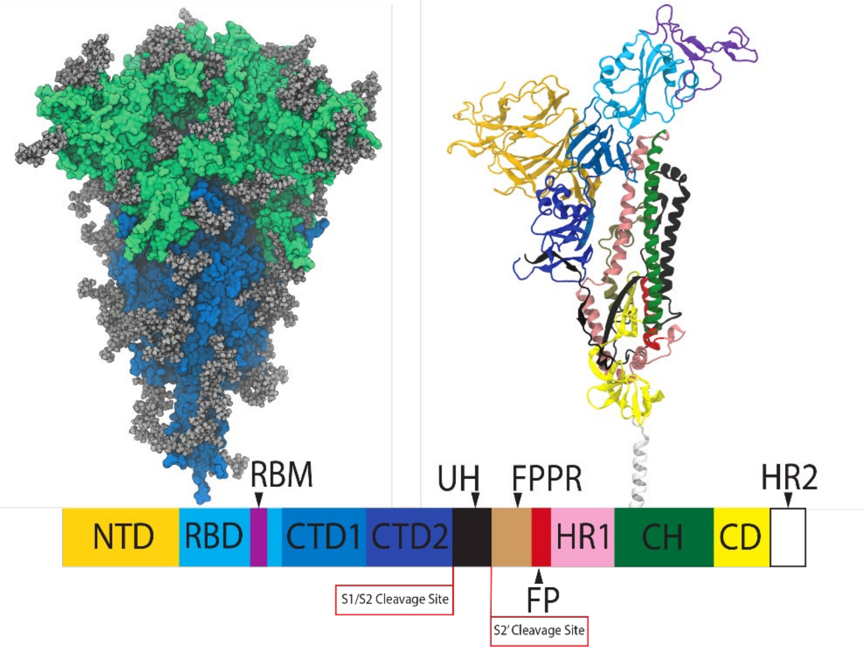


**Figure S1.** (A, upper left) structure of the trimeric SARS-CoV-2 spike glycoprotein ectodomain, with the S1 subunit shown in green, the S2 subunit shown in blue, and the glycans in gray. (B, right) Ribbon drawing of a single protomer, color coded by domains indicated in the lower figure. Domain labels are adapted from prior work(1).

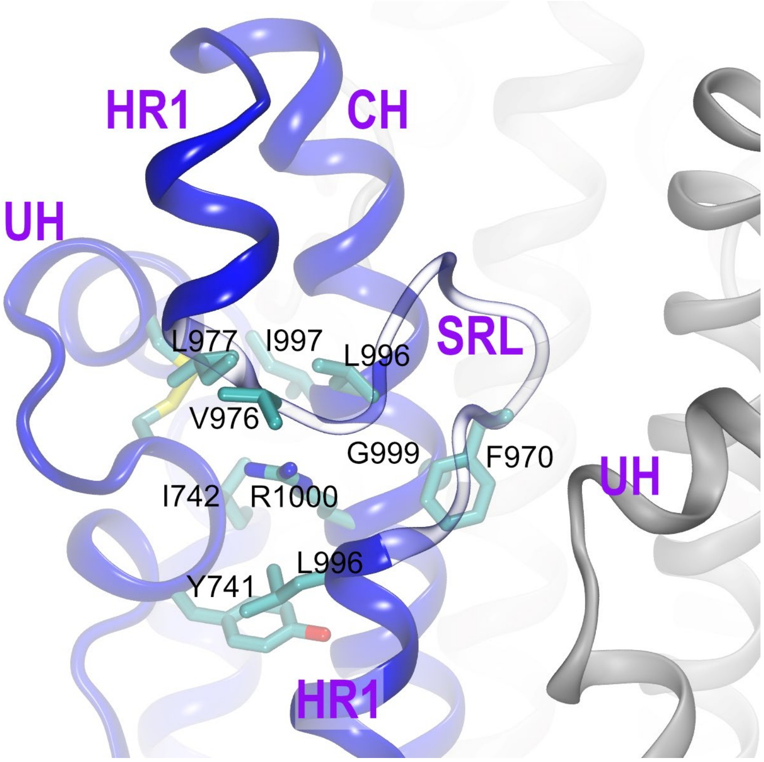


**Figure S2**: The side chain of R1000 on the CH is surrounded by hydrophobic amino acids. The phenyl ring of F970 on the SRL is packed against the α-carbon of G999 on the CH.  The structure shown is the wild-type spike in the pre-fusion conformation (6XR8). The SRL (shown transparent to aid visualization) forms a cap over the hydrophobic pocket.

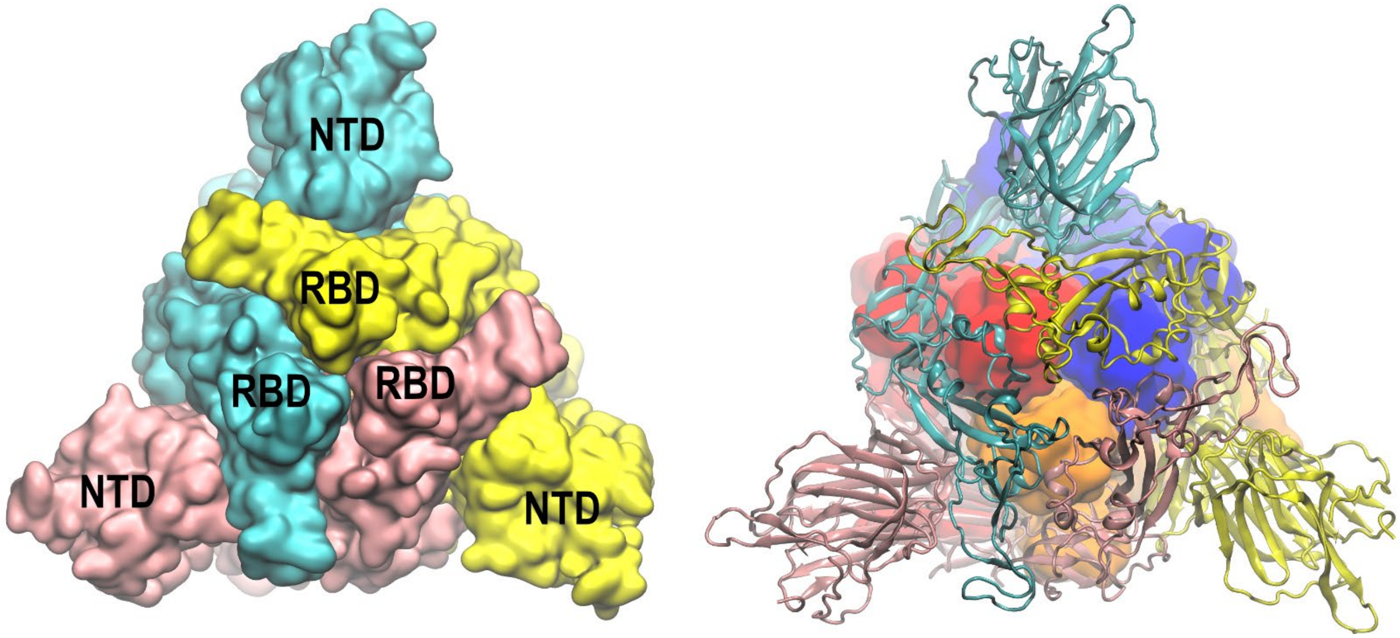


**Figure S3.** Top view of the pre-fusion spike 6XR8, showing locations of NTD and RBD domains of the S1 subunit atop the spike. **Left**: Each protomer is shown in a different color space-filling model (red, blue and yellow in a clockwise rotation). **Right**: The S1 subunit is shown in ribbon drawing, while the lower S2 subunit is shown as space-filling. The S2 protomers are shown in a darker shade than the same protomer in the S1 subunit. Each RBD covers the S2 subunit of both of the other protomers; correspondingly, each **S2 subunit is covered by the RBD domains of both of the other protomers**.

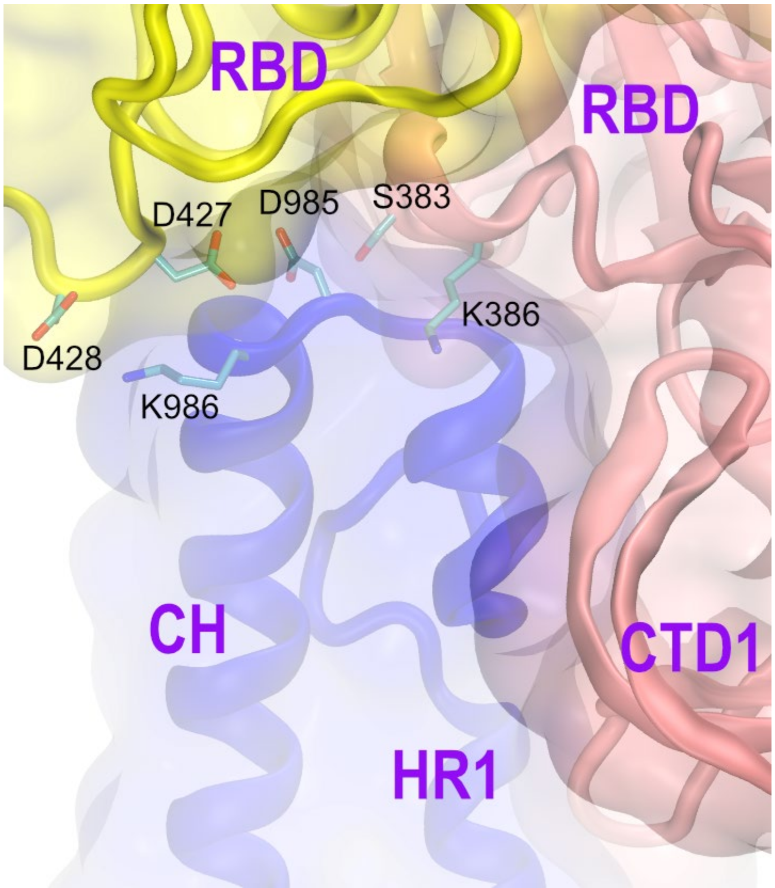


**Figure S4.** The RBD domains of 2 different protomers cover the helix-turn-helix involving CH and HR1 at the top of one S2 protomer. Ribbons are colored by protomer. Labeled amino acids are discussed in the main text. The structure shown is the wild-type spike in the pre-fusion conformation (6XR8).

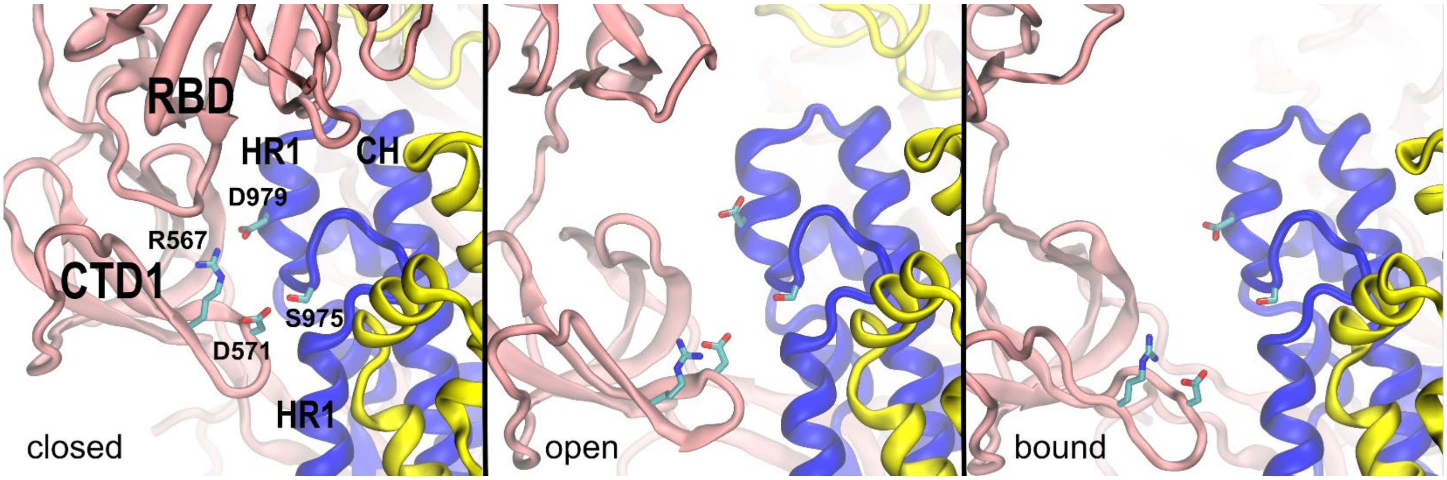


**Figure S5.** Contacts between the CTD1 and the HR1 upper short helix, in 3 cryo-EM structures of the pre-fusion spike. Ribbons are colored by protomer. Left: 6XR8, closed. Middle: 6VSB, open unbound. Right: 7CAK, 3 RBDs open and bound. As the RBD opens, the distance between CTD1 and HR1 increases, resulting in loss of contacts between the S1 and S2 subunits (D571-S975 and R567-D979).

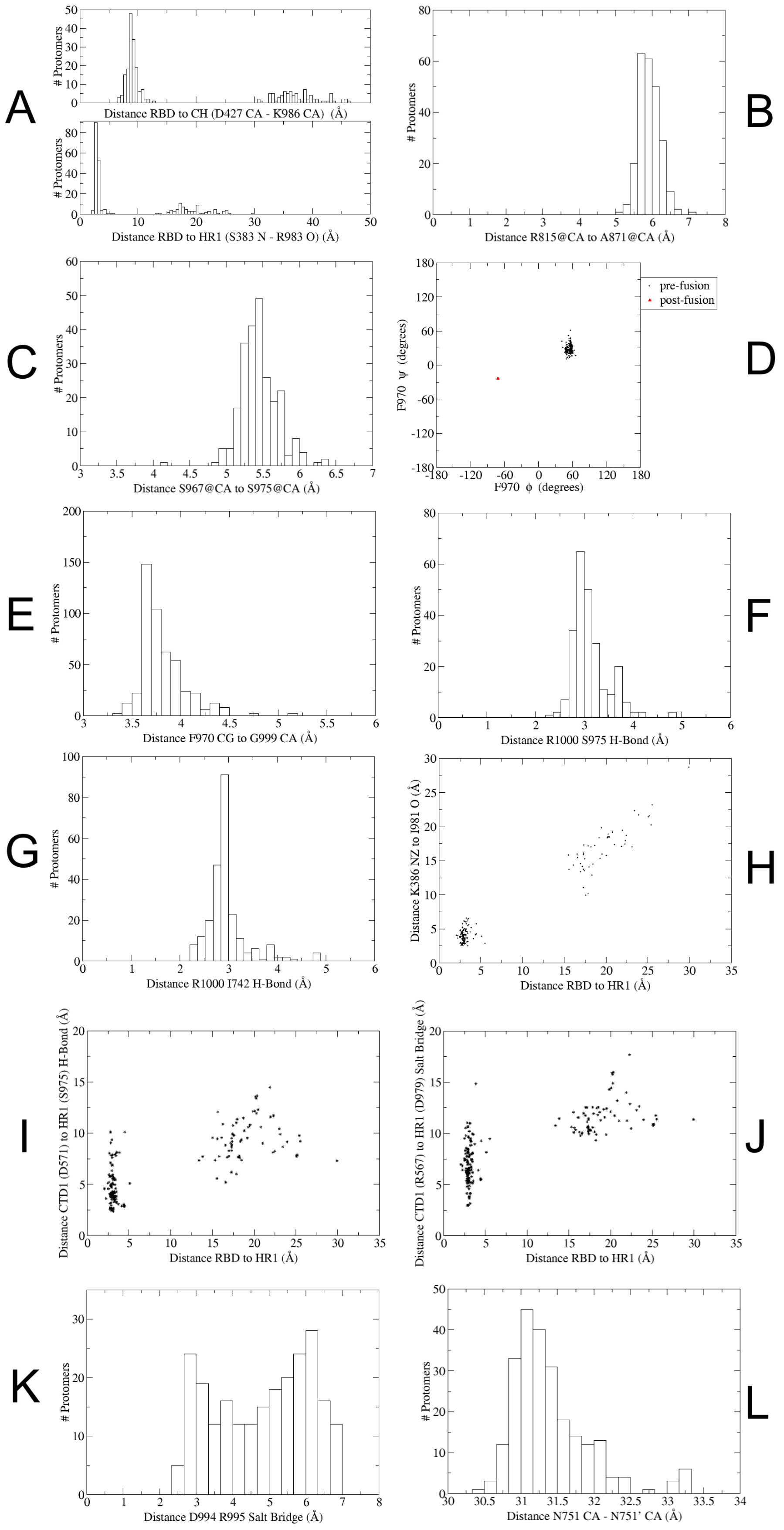
A
### Protomers
Distance RBD to CH (D427 CA - K986 CA) (Å)
Distance RBD to HR1 (S383 N - R983 O) (Å)
B
Distance R815@CA to A871@CA (Å)
C
Distance S967@CA to S975@CA (Å)
D
F970 ψ (degrees)
F970 ϕ (degrees)
pre-fusion
post-fusion
E
Distance F970 CG to G999 CA (Å)
F
Distance R1000 S975 H-Bond (Å)
G
Distance R1000 I742 H-Bond (Å)
H
Distance K386 NZ to I981 O (Å)
Distance RBD to HR1 (Å)
I
Distance CTD1 (D571) to HR1 (S975) H-Bond (Å)
J
Distance CTD1 (R567) to HR1 (D979) Salt Bridge (Å)
K
Distance D994 R995 Salt Bridge (Å)
L
Distance N751 CA - N751' CA (Å)

**Figure S6.** Data extracted from 80 SARS-CoV-2 spike pre-fusion structures obtained from the PDB (PDB codes provided in **Table S1**). (**A**) Histogram of distances from RBD to the top of HR1 or CH (the leftmost peak corresponds to closed RBDs). (**B**) Histogram of distances from R815 at the S2' site to nearby A871. (**C**) Histogram of distances for the close approach of the ends of the SRL, from S967 to S975. (**D**) Backbone Ramachandran plot for F970, showing that all pre-fusion structures (black) adopt a high-energy region of the phi/psi space, in contrast to the post-fusion 6XRA (red). (**E**) Histogram of distances from the F970 phenyl ring to G999. (**F**) Histogram of distances for the hydrogen bond from R1000 side chain to the S975 backbone O. (**G**) Histogram of distances for the hydrogen bond from R1000 side chain to the I742 backbone O. (**H**) X-axis, distance from RBD to the top of HR1 (S383 N to R983 O); Y-axis, C-capping distance from K386 NZ (RBD) to I981 O (HR1) (see **Figure S4**). (**I**) X-axis, distance from RBD to the top of HR1 (S383 N to R983 O); Y-axis, hydrogen bonding distance from D571 (CTD1) to S975 (HR1) (see **Figure S5**). (**J**) X-axis, distance from RBD to the top of HR1 (S383 N to R983 O); Y-axis, salt bridge distance from R567 (CTD1) to D979 (HR1) (see **Figure S5**). (**K**) Histogram of distances for possible salt bridge between D994 (CH) and R995' (CH') (**Figure S7**). (**L**) histogram of distances corresponding to edges of the UH-UH triangle (**Figure 3**): distance from N751 CA to N751' CA.

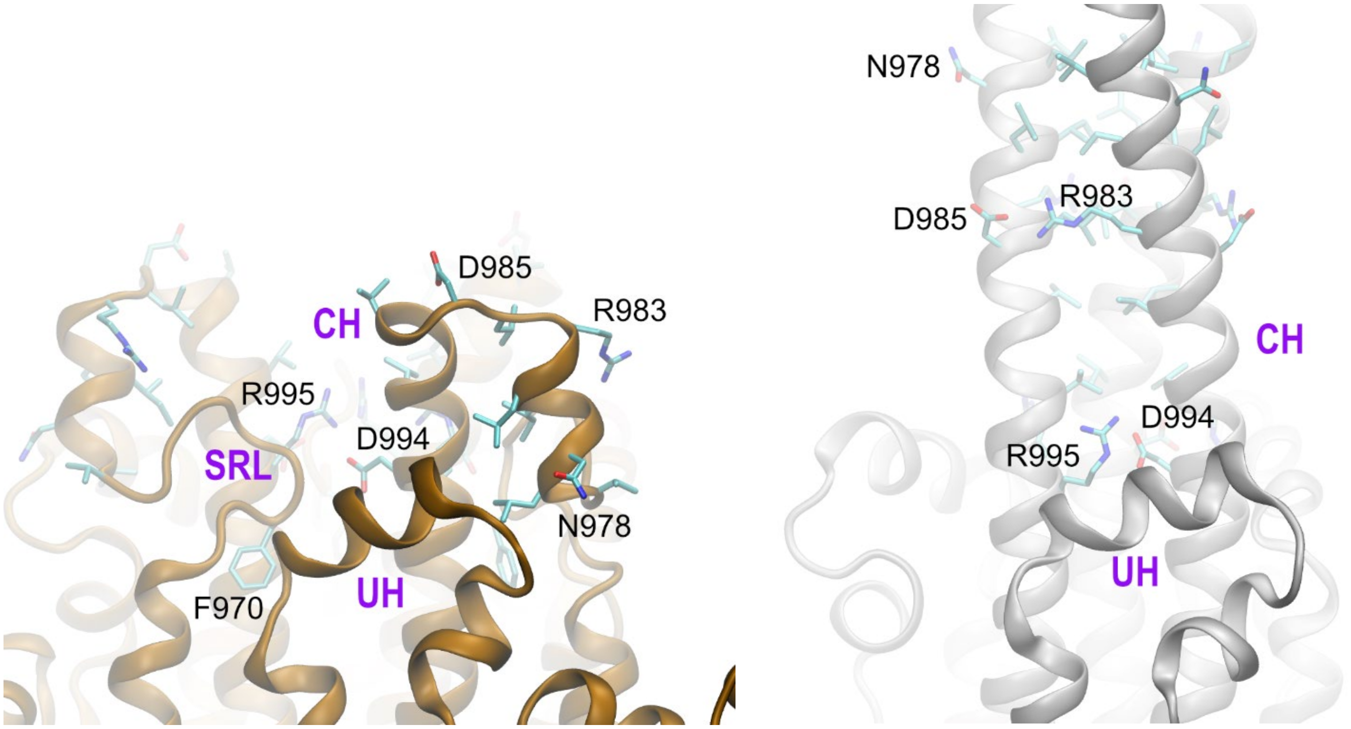


**Figure S7**. View of selected amino acids at the top of the wild-type spike S2 subunit in the pre-fusion (left, 6XR8, copper) and post-fusion (right, 6XRA, silver) structures. In the post-fusion spike, six new salt bridges have formed, separated by only 3 turns of α-helix. These involve D994/R995' and R983/D985' on the CH of neighboring protomers. A new hydrophobic cluster stabilizes the post-fusion coiled-coil motif above R983.

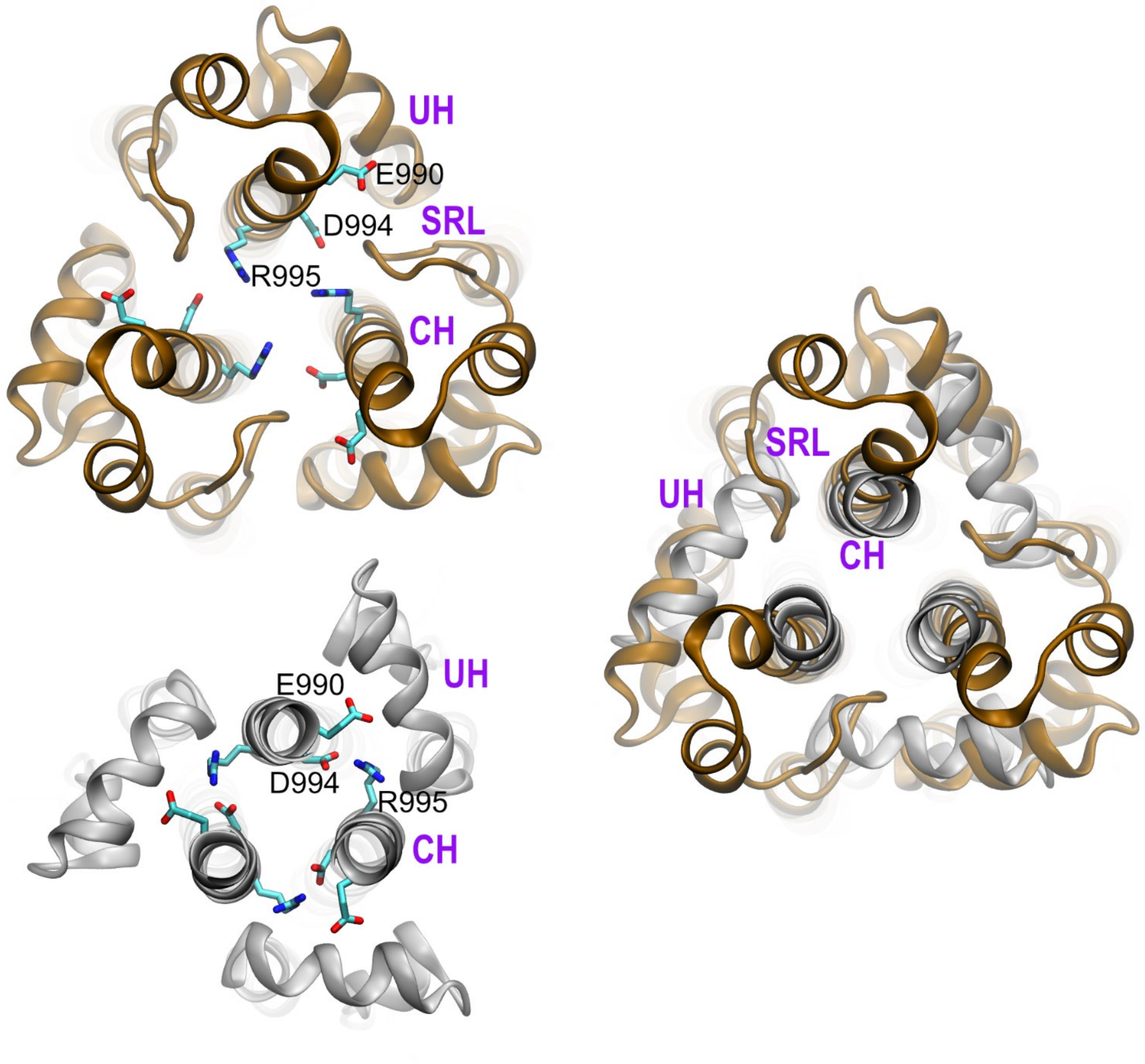


**Figure S8**. Cross-section of the wild-type spike near the SRL, viewed from the top. (Top left) pre-fusion (6XR8, copper) and (bottom left) post-fusion (6XRA, silver), and (right) both structures best-fit to CH. Labels are used on selected amino acids for one protomer. In the pre-fusion structure, the SRL inserts between the CH and UH of neighboring protomers. In the post-fusion structure, SRL is relocated, the CH and neighbor UH are in contact, and salt bridges D994-R995’ form between the CH of neighboring protomers. Overlap indicates that the SRL blocks the closer approach of CH and neighbor UH in the pre-fusion structure.

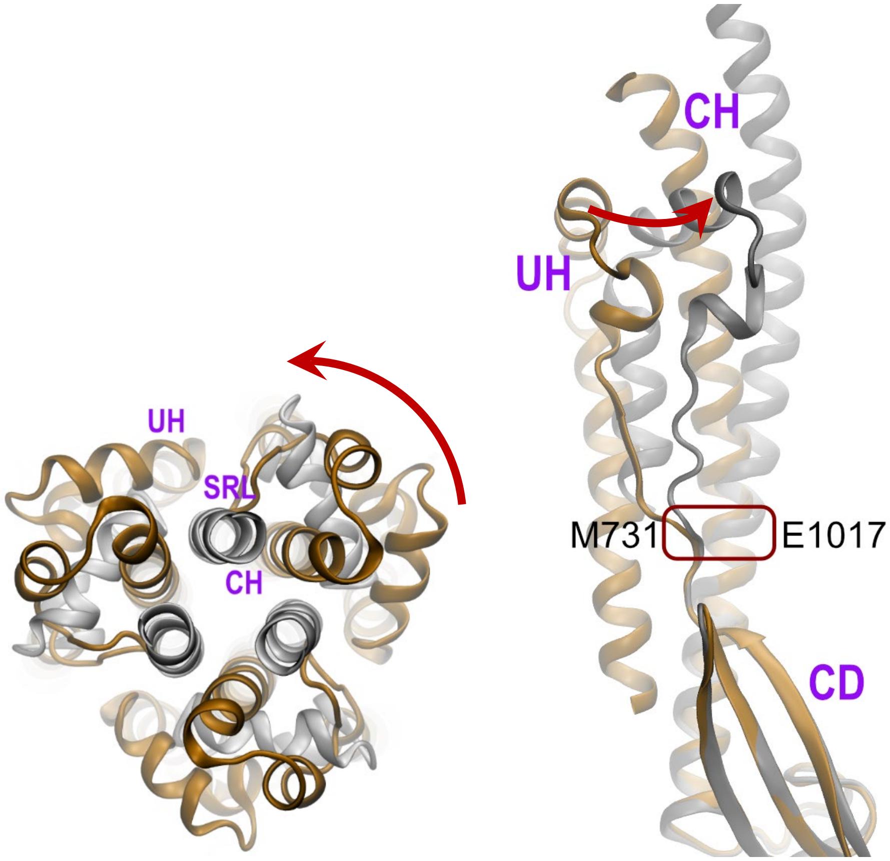


**Figure S9**. The upper S2 rotates counter-clockwise relative to the lower S2 during spike activation (indicated by red arrows). Top view (left) and side view (right) of the wild-type spike in pre-fusion (copper, 6XR8) and post-fusion (silver, 6XRA) structures. The structures are best-fit to the lower part of the spike ($L_{1049}MSFPQSAP_{1057}$) in the connector domain (CD). The fulcrum for rotation of the upper S2 is indicated by the red box drawn at M731 for UH, and E1017 for CH. For clarity, some regions are omitted from view.

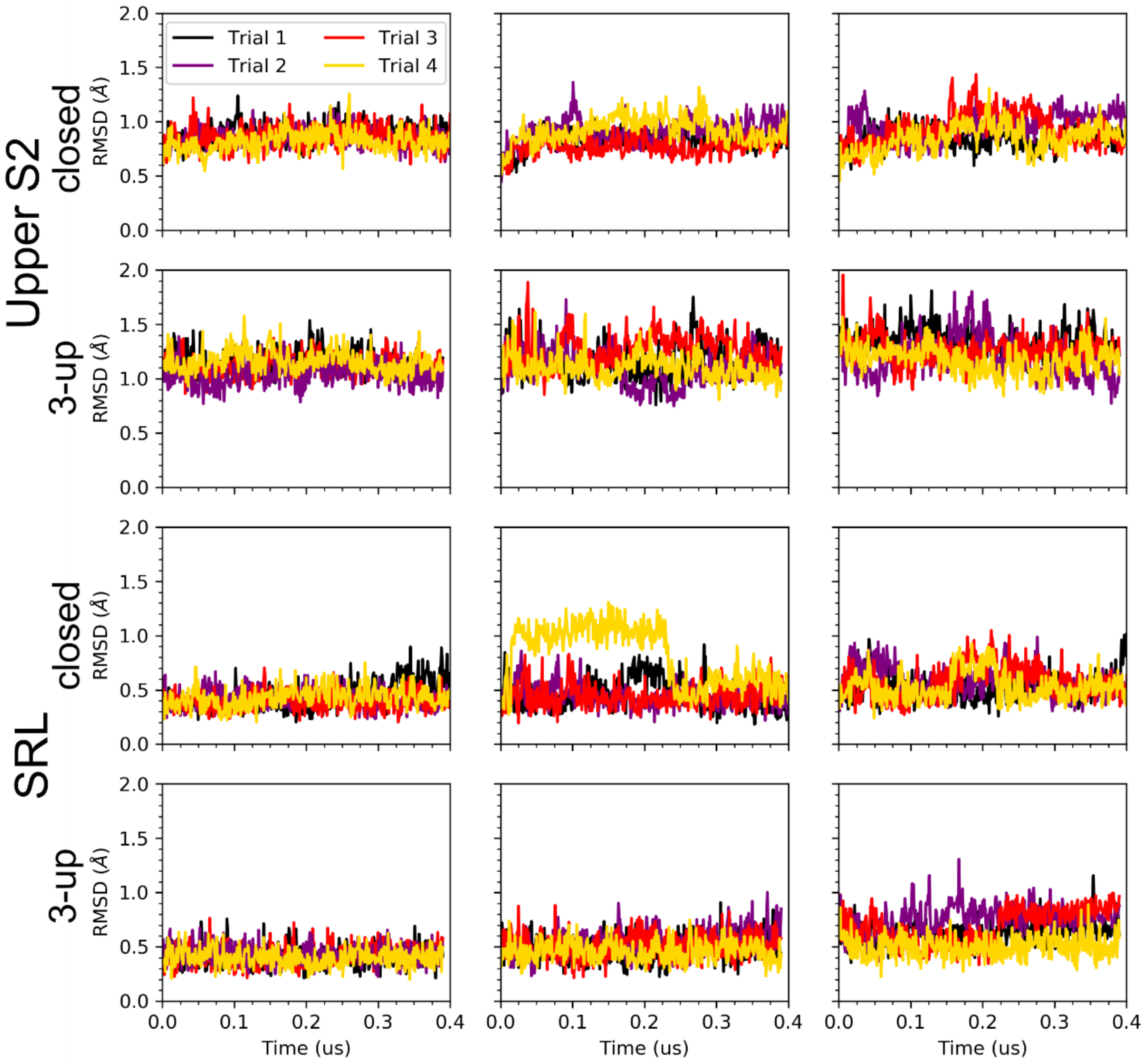


**Figure S10.** RMSD analysis of the closed and 3-up standard MD simulations of the full spike ectodomain. Each column represents a different protomer. The different colors on each plot represent four independent trials of ~ 400 ns each that were carried out for closed and 3-up spike. The upper S2 region includes positions 737-761 (UH) and 962-1005 (HR1-SRL-CH). The SRL region includes positions 967-976. The reference structure for all plots was the 6XR8 closed structure. The S2 and SRL are stable in all runs.

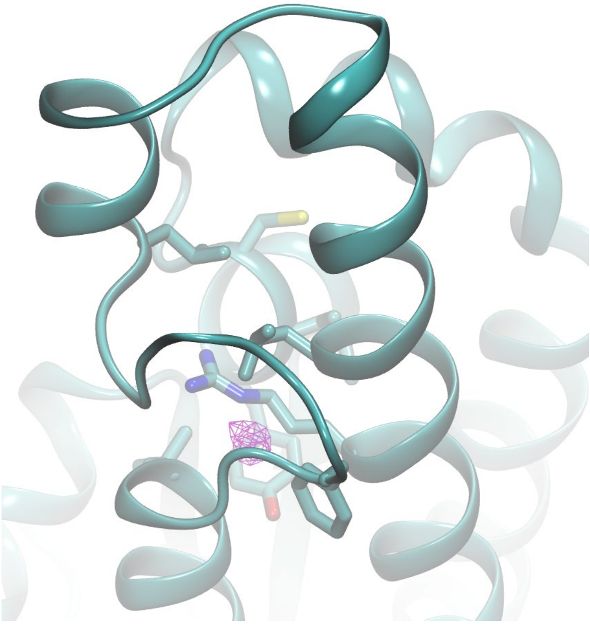

**Figure S11.** Water density grid (purple) inside the hydrophobic pocket that encloses R1000, calculated during MD simulation of the closed full spike ectodomain. Similar water occupancy at this location is observed in all MD simulations.

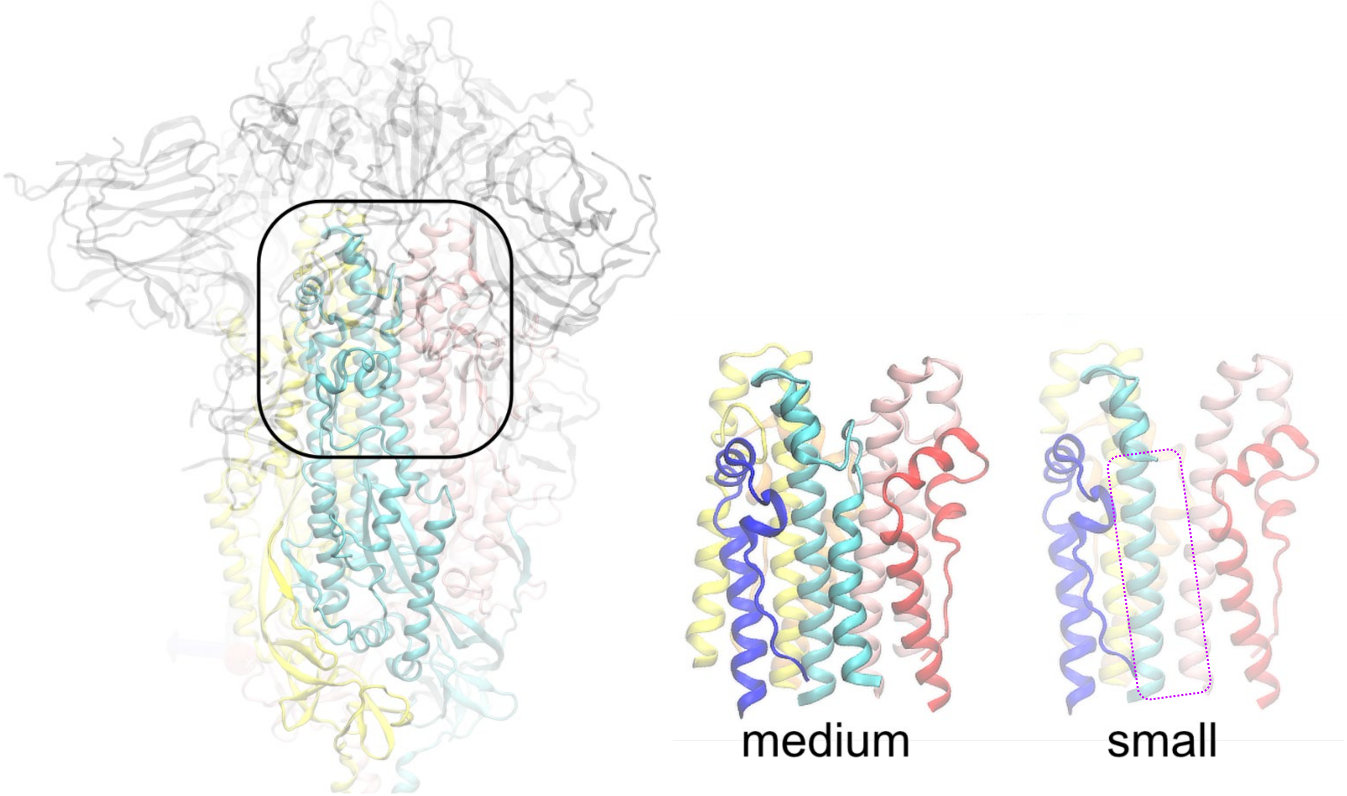


**Figure S12**: Two model systems that were used for simulations of the upper region of the S2 subunit. Left: the pre-fusion spike trimer 6XR8, with the S1 subunit in gray and S2 colored by protomer. The box indicates the area corresponding to the model systems. Middle: the medium model system, retaining only the CH, HR1, SRL and UH, truncated at the bottom. Right: the small model system obtained by removing the SRL and lower helix of HR1 from the medium model (indicated by the dashed box).

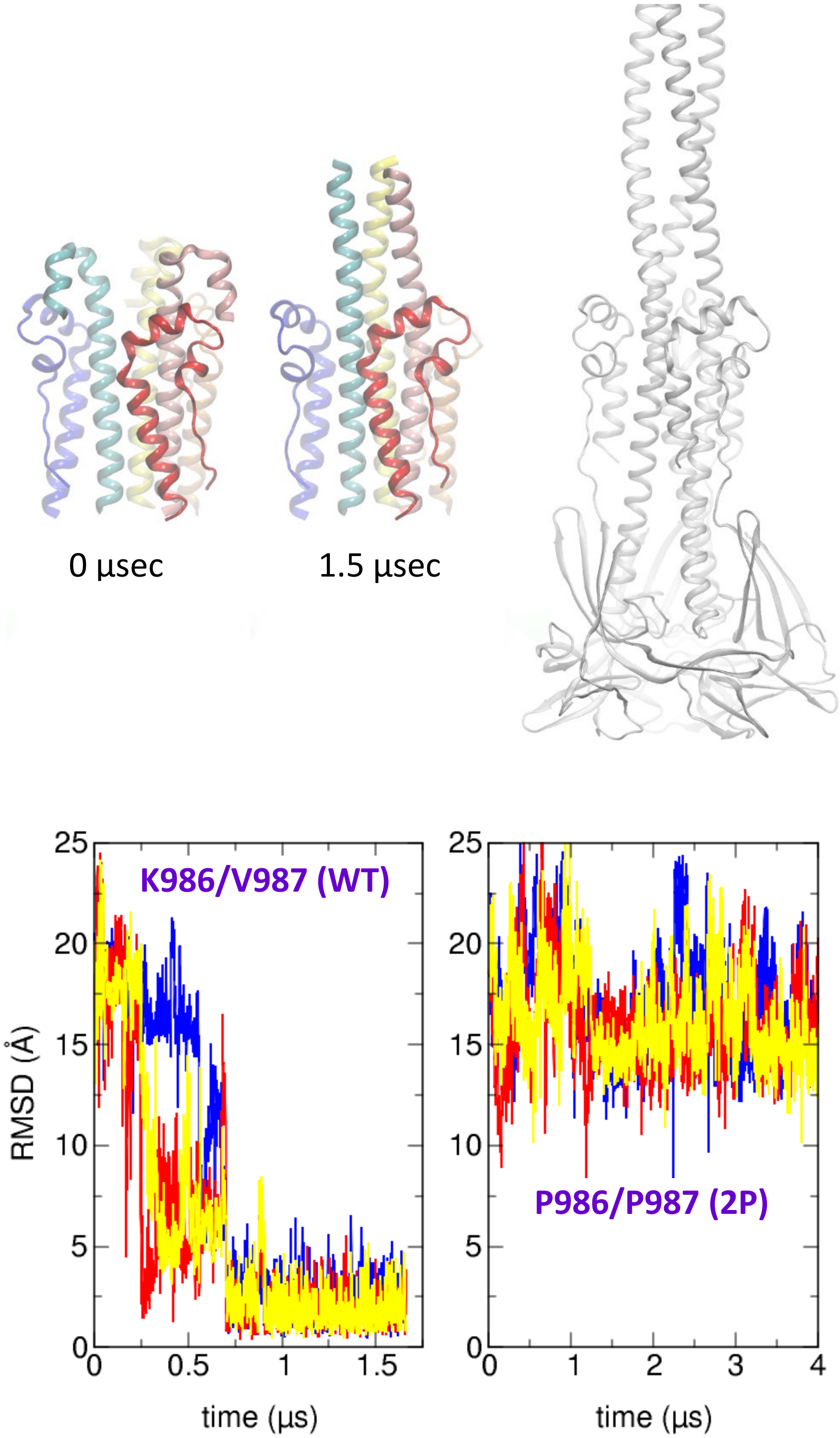


**Figure S13**: Results of standard all-atom MD in explicit water on the small model system compared to the post-fusion spike 6XRA. (**Upper**) Small model structure (primary colors, UH in darker shade) and post-fusion spike (gray, right), all best-fit to amino acids in the CH near the base of the model system ($Q_{1005}TYVTQQLIRAA_{1016}$). The initial conformation (left, initiated from the pre-fusion structure 6XR8) differs mainly in the helix-turn-helix motif at the top of the CH. After 1.5 µsec standard MD (middle), the upper short helix segments of HR1 have spontaneously rotated upward and lengthened the CH of each protomer. (**Lower)** Time-dependent backbone RMSD values of each protomer for the upper HR1 helix-4 segment ($D_{979}ILSRL_{984}$) in simulations compared to the post-fusion structure (after best-fit to CH $K_{986}VEAEVQIDRLITG_{999}$). Left: wild-type sequence; right: including 2P substitution, in longer simulations. **The HR1 rotates to extend the CH in the wild-type, but not in the 2P system.**

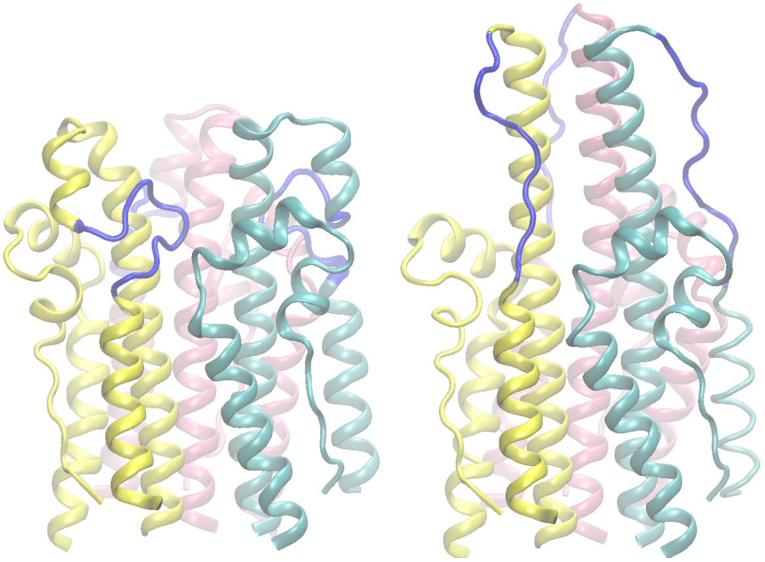

**Figure S14**. Unfolding of the SRL allows extension of the central helix. Medium model system before (left) and after (right) extension of the CH by using SMD on the HR1 helix-4 to match the structure obtained in the small model MD (**Figure S13**). Protomers are shown in light primary colors, and the SRL is shown in dark blue.

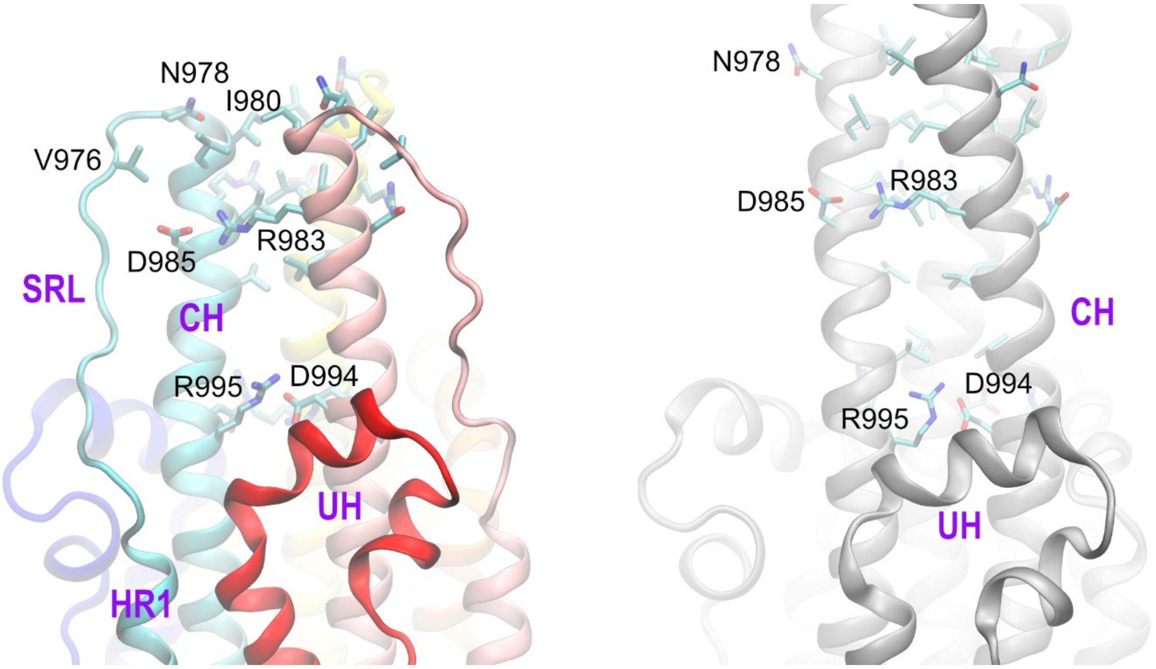


**Figure S15**. Comparison of the final structure from MD on the S2 medium model to the post-fusion spike. (Left) MD structure after CH extension, with each protomer in a different primary color (CH/HR1 lighter, UH darker). N978 forms an N-capping interaction on the CH, similar to that involving D985 in the pre-fusion structure (**Figure 2**). Other contacts that form are comparable to the post-fusion structure (right image), including six additional salt bridges between neighboring CH pairs, and a hydrophobic cluster involving I980 and L984 that stabilizes the central coiled-coil.

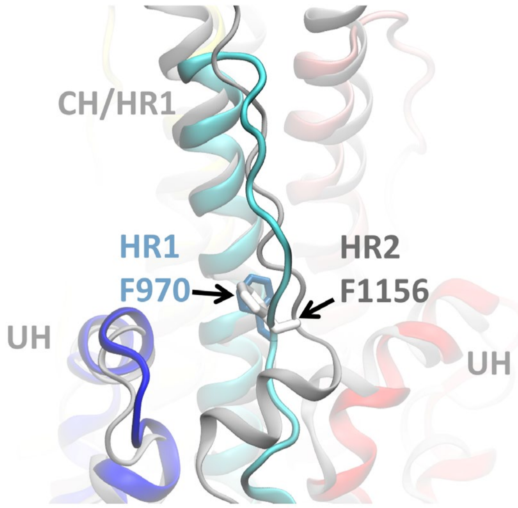


**Figure S16**. Comparison of the medium model system after MD (primary colors) with the post-fusion structure 6XRA (gray). The unfolded SRL follows a similar path alongside CH to that taken by a segment of HR2 in the post-fusion structure. The side chain of F970 is positioned similarly to F1156 on HR2 in the post-fusion structure. For clarity, some regions are omitted.

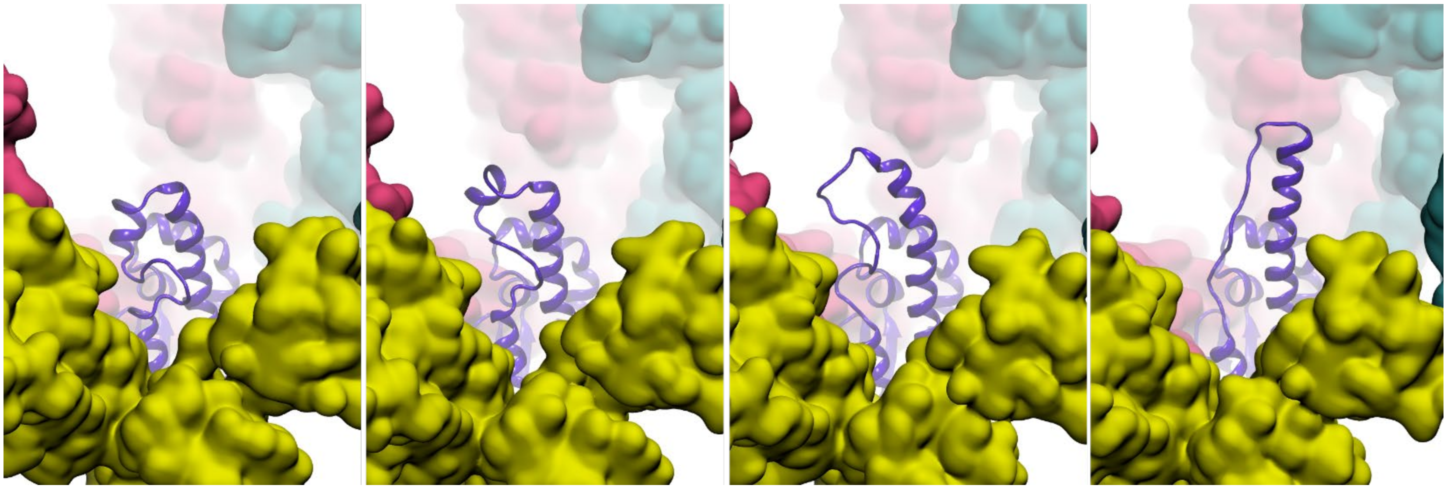

**Figure S17.** Snapshots along the pathway for RBD extension in the 3-up full spike ectodomain system (with the initial pre-fusion structure shown on the left). A single protomer of S2 is shown using purple ribbons, and the S1 subunits are shown in space-filling with a different color for each protomer. **For clarity, only a single protomer of S2 is shown**, and the RBD of the yellow S1 subunit is omitted to provide a clear view of the S2.

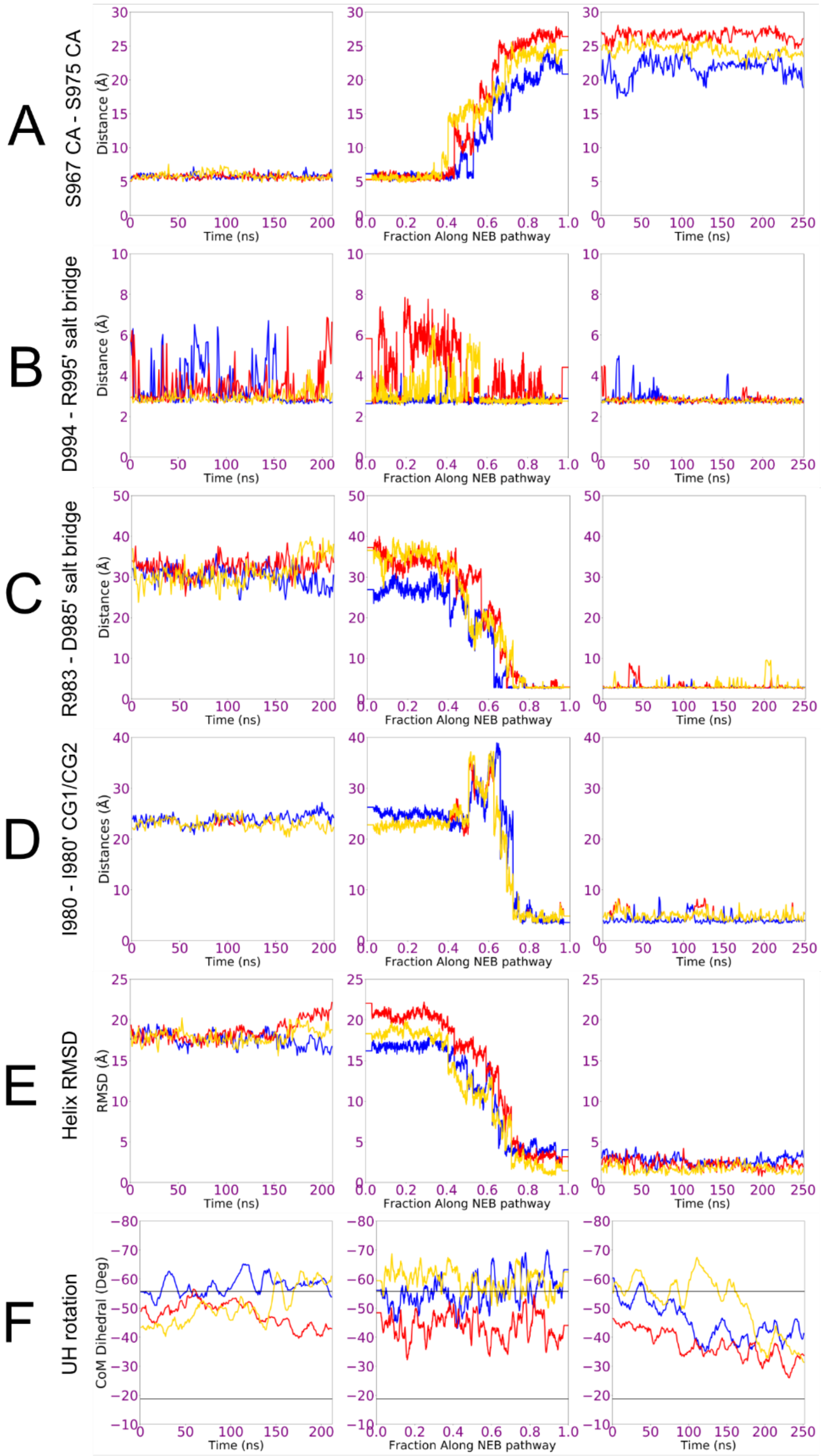


**Figure S18**. Time dependence of changes in full spike ectodomain system during the three steps of (left column) unrestrained MD for 3-up spike; (middle column) SMD for extension of CH using HR1 helix-4; (right column) unrestrained MD of 3-up spike after SMD. Each protomer is shown in a different color. (A) Distance between CA atoms of S967 and S975 of the SRL that approach closely in the pre-fusion state (structure in **Figure 2**); (B) salt bridge distance between D994 (CH) and R995' (CH) on the neighboring protomer (structure in **Figure S15**); (C) salt bridge distance between R983 (HR1) and D985' (CH) of the neighboring protomer (structure in **Figure S15**); (D) distance between CG atoms of I980

(HR1) and I980' of the neighboring protomer (structure in **Figure S15**), (E) backbone RMSD of the upper HR1 segment ($D_{979}ILSRL_{984}$) in simulations compared to the post-fusion structure (as shown in **Figure S13** for the model system), (F) twist of the UH in each protomer (structure in **Figure S9**). Upper and lower lines indicate values from the pre-fusion and post-fusion cryo-EM structures, respectively. *After partial extension of CH, the newly formed contacts are stable during 250 ns of unrestrained MD, and the UH slowly relax toward the value adopted in the post-fusion structure.*

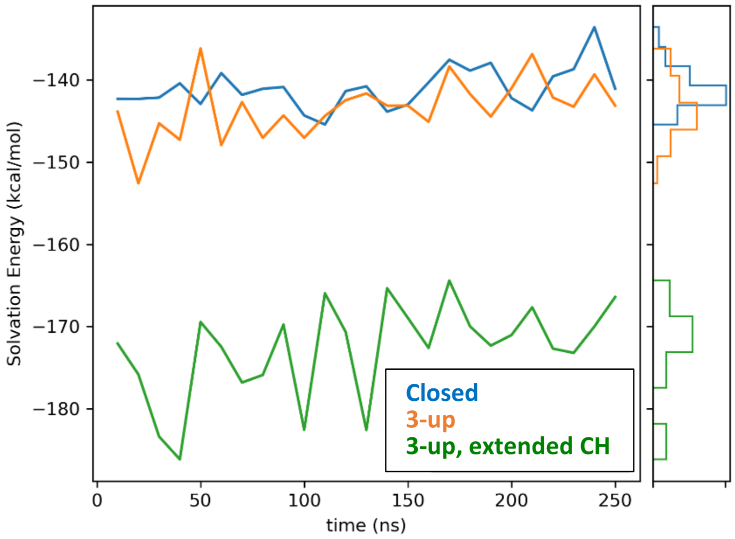


**Figure S19.** Combined solvation free energy of the three R1000 amino acids, during MD trajectories of different spike conformations. Calculations were performed on MD snapshots using the Poisson-Boltzmann method. Blue: closed; orange: 3-up, green: 3-up, partially extended CH. The extension of CH and unfolding of the SRL leads to significantly better solvation energy for R1000.

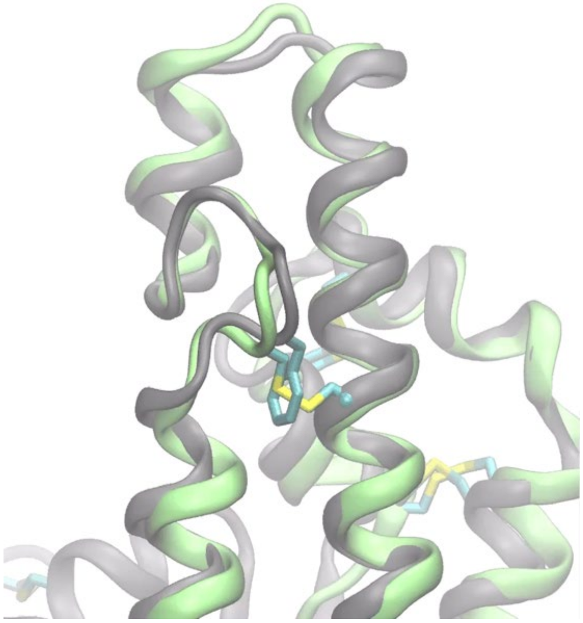

**Figure S20**: Overlap of one protomer of S2 from: (green) a full-spike ectodomain MD simulation with the F970C/G999C substitution and disulfide bond, and (gray) the pre-fusion structure 6XR8 (F970/G999). Disulfide bonds and F970/G999 are shown in licorice. The introduction of the disulfide is accommodated in simulations with minimal perturbation. For clarity, other protomers and the S1 subunit are not shown.

**Table S1. PDB codes for the 81 experimental spike structures used in data analysis. All structures are pre-fusion except 6XRA.**

6VSB, 6VXX, 6WPS, 6WPT, 6X29, 6X2A, 6X2B, 6X2C, 6X6P, 6XCM, 6XCN, 6XEY, 6XF5, 6XKL, 6XLU, 6XM0, 6XM3, 6XM4, 6XM5, 6XR8, 6XRA, 6Z43, 6Z97, 6ZDH, 6ZGE, 6ZGG, 6ZGH, 6ZGI, 6ZOW, 6ZP5, 6ZP7, 6ZWV, 6ZXN, 7A29, 7A4N, 7A93, 7A94, 7A95, 7A96, 7A97, 7A98, 7AD1, 7BYR, 7C2L, 7CAI, 7CAK, 7CHH, 7CN9, 7JJI, 7JV4, 7JV6, 7JVC, 7JW0, 7JWB, 7JZL, 7JZN, 7K43, 7K4N, 7K8S, 7K8T, 7K8U, 7K8V, 7K8W, 7K8X, 7K8Y, 7K8Z, 7K90, 7KDG, 7KDH, 7KDI, 7KDJ, 7KDK, 7KDL, 7KE4, 7KE6, 7KE7, 7KE8, 7KE9, 7KEA, 7KEB, 7KEC

#### Additional Methods Details

*Building the spike model with all RBDs closed*

Our model of the SARS-Cov-2 spike protein ranged from residue Q14 to V1164, covering most of the parts of S1 and S2. To avoid the influence of charged termini, we introduced three acetyl capping groups to the 3 N-terminals of the 3 protomers. Model building started from the closed model for the glycosylated spike that we described previously(2), with additional changes described here in order to incorporate additional structural details that are resolved in newer cryo-EM structures. Steered MD (SMD) was used to change the structure to match the wild-type closed spike in cryo-EM structure 6XR8, which resolved the coordinates for the fusion peptide proximal region FPPR(1) under the CTD12 domain; this is missing in many other structures. The collective variable (CV) used for SMD was RMSD with all CA atoms in 6XR8. The RMSD was gradually reduced from the initial value of 5.2 Å to 0, using 100000 kcal/(mol Å$^2$) harmonic restraint during 20 ns. After SMD, a few MD and SMD steps were applied to fix some glitches in the model introduced during SMD. We did additional equilibration on the sidechains and glycans that were not included in the SMD restraint. While keeping the same CA RMSD restraint of 100000 kcal/(mol Å$^2$), the system was heated from 310 K to 400 K during 2 ns, kept at 400 K for 16 ns and gradually cooled back to 310 K. The RMSD restraints were released and the system was again heated to 400K during 2 ns, then kept at 400K for 8 ns. A loop region not resolved in 6XR8 was observed to be distorted, so we added another CV on a loop region of S1 (residue A61 to F79) using the CA atoms of the same loop of protomer 2 from the first heating step as reference, and SMD was applied at 400 K over 20 ns while maintaining the RMSD restraint. A glycan branch (connected to N17 of protomer 1 in our model) was found to have become trapped between protein chains; it was removed with additional SMD on two atoms. The distance between the C1 atom of residue 689 of our model, which is in the glycan branch connected to N74 of protomer 1, and the CA of Glu339 of protomer 1 RBD was increased from 50.5 Å to 110 Å during 14 ns of SMD at 400 K. No RMSD based restraint was used in this step. The SMD restraint using 6XR8 as reference was added back. The total RMSD was reduced from 1.26 Å to 0.21 Å by gradually increasing the force constant from 1000 to 100000 kcal/(mol Å$^2$) during 10 ns. The simulation temperature was gradually decreased during the first 8 ns from 400 K to 310 K, then kept at 310 K for the last 2 ns. Another glycan problem was fixed in the same way. Atoms C1 of residue 1997 of our numbering, which is in the glycan branch connected to N17 of protomer 2 of our model, and atom CA of C136 of protomer 2 of our model was selected for SMD, and their distance was increased from 12.2 Å to 30 Å during 4 ns at 310 K. Next, we adjusted the chemical composition of our spike model to better match 6XR8. Short sequences Ace-Asn14-Cys15 were added to the 3 N-termini of S1 to resolve a missing disulfide bond. Disulfide bonds were added between C15 - C136 (in NTD) and C840 – C851 (in FPPR). All O-linked glycans were deleted from the original model due to doubts(3, 4) about their presence in the trimer. The spike protein was solvated using 31 Å minimum distance from solute to box edge with 371793 water molecules added. A large box was used to ensure adequate solvation of the flexible glycans, and enclosure of the RBDs after opening. 373 sodium ions and 353 chloride ions were added. The same equilibration protocol was used as our previous work(2), except that in the first minimization step the default 10 steps of steepest descent algorithm were used followed by conjugate gradient. 6XM5 also resolved a loop region in CTD2 missing from our original model template. The CA atoms of V608 to G652 of chain A from 6XM5 were used as reference. The RMSD of three regions were reduced from their initial value (around 5-6 Å) to 0 using 50000 kcal/(mol Å$^2$) restraint in 10 ns SMD. Finally, the FPPR structure solved in PDB 6XM5(5) was used as an alternative FPPR conformation (hereafter referred as "Kwong FPPR", with the one from 6XR8 referred as "Chen FPPR"). SMD was used to change the conformations. CA atoms of S816 to D867 of chain B of 6XM5 were used as the structural reference. The RMSD of the three FPPRs was reduced from around 8 Å to 0 using 50000 kcal/(mol Å$^2$) restraint during 10 ns SMD. Separate structures were built using both the Chen FPPR and Kwong FPPR; the Chen FPPR was employed for closed states (consistent with 6XR8), and the structure with Kwong FPPR was used for building the 3-up models since the Chen FPPR creates steric clash with the CTD1 location in 7CAK.

*Building the spike model with all RBDs open (3-up)*

We built our 3-up model based on the cryo-EM model of the 3-up spike with each RBD bound to an H014 antibody Fab fragment in the 7CAK structure. We began with the closed spike model. All 3 RBDs were opened by SMD using our equilibrated 1-up model from our previous work(2) as reference. The RMSD selection included CA atoms of residues 338-517,324-327,538-585,747-782,946-966 and 987-1034, which include the RBD, CTD1 and helical segments in UH, HR1 and CH. 3 RMSDs was reduced from ~ 12.5 Å to 0 using 100,000 kcal/(mol Å$^2$) restraint over 20 ns. After generating the 3-up model, 270 ns of regular MD was carried out. Since differences were noted in the experimental structures of the 1-up and 3-up spike, the 3-up structure after MD was steered to match 7CAK by reducing RMSD from 12.2 Å to 0 using 100,000 kcal/(mol Å$^2$) force constant over 30 ns. All CA atoms resolved in 7CAK were used for steering. The RMSD restraint was maintained on the CA atoms for an additional 30 ns to relax the model before unrestrained MD runs. All simulations were run at 310 K.

*Building the model systems with a subset of the S2 trimer*

Two small model systems for the S2 subunit were built, both based on the wild-type pre-fusion 6XR8 structure. This approach is similar to the peptide fragment experiments used to determine the spring-loaded mechanism of influenza hemagglutinin(6, 7). All amino acids below the fulcrum of S2 rotation **(Figure S9)** were removed. In the "medium" system, we retained the central components of all three protomers in the trimer, including the CH, SRL, HR1 and UH (M731-K776, D950-R1019) (**Figure S10**). The N- and C-terminal ends of all six chains were capped with neutral termini and restrained to their initial coordinates during all MD steps described below. In the "small" model, we further truncated the medium system by deleting the HR1 chains from D950 to S975, with neutral capping added but no restraints applied to V976. Both systems were simulated in explicit water.

The small model system was initially simulated in TIP3P 3-point water model for speed, since our goal was to carry out a long unrestrained simulation to test for helix extension. We used the ff14SB force field that is optimized for TIP3P. Water was added with a minimum distance of 8 Å from solute to the box edge, resulting in addition of 6054 water molecules. The system was equilibrated in a series of stages including 1000 steps of minimization, 100 ps of heating from 100 K to 298K at constant volume and with positional restraints of 10 kcal/mol/Å$^2$ on all non-solvent atoms; 200 ps at constant pressure to equilibrate density with positional restraints of 10 kcal/mol/Å$^2$ on non-solvent atoms; 200 ps with positional restraints of 10 kcal/mol/Å$^2$ on the capping groups at the base of the fragment (see above). This was followed by 1.65 microseconds of MD at 350K and 1 bar, maintaining the restraints on the fragment caps. The increased temperature was employed based on experiments(6) that show elevated temperature leads to activation of hemagglutinin at neutral pH. Refinement was carried out for the final structure by converting to the more accurate but slower ff19SB force field with OPC 4-point explicit water. The structure was equilibrated using the same protocol as described above, followed by 1 microsecond of MD at 310K with restraints only on the lower fragment caps.

The medium model system simulations aimed to create a model for the unfolded SRL and location of the new, longer CH α-helix. We retained the ff19SB force field with the OPC water model. The system was equilibrated using the same protocol as the small model system. Following 400 ns of MD at 330K and 1 bar, the final snapshot was used to initiate SMD simulations to extend the central helices. The medium model was steered to unfold the SRL and extend the CH cap by using the uncapped small model as reference. A separate CV was applied to each protomer. The RMSD region included CA atoms of N978 to E1017 (including the short upper HR1 helix, continuing down CH). N978 was chosen as the end of the restrained region since the simplest hypothesis for extension of CH was that the upper 2-turn α-helix of HR1 (L977-L984) would add to CH, and that the SRL (S967-V966) would remain non-helical.

As with the small model, the free ends of the protein chains at the base of the model were restrained to their initial positions using 10 kcal/(mol Å$^2$) force constant. RMSDs for the CV regions were reduced from around 6 Å to 0 using 10000 kcal/(mol Å$^2$) restraint during 40 ns at 330 K. Following SMD, the structure was relaxed for 110 ns at 330 K to optimize the conformation of the SRL linking the extended CH and the lower HR1. Distance restraints were applied to maintain hydrogen bonds on the newly-extended helix, and positional restraints on the fragment base were maintained. Next, a simulation of 887 ns was carried out for the same system at 300 K and 1 bar, again using FF19SB and OPC, with restraints only on the fragment base. The final snapshot was used as an SMD target for the full spike system.

*Building the 3-up spike with partially extended CH and unfolded SRL*

The model building started from the MD snapshot of the 3-up model after 210 ns unrestrained MD. The final structure of our medium model with all 3 CH extended (**Figure S14**) was used as a structural reference. The RMSD region included the CA atoms of residues K964 to Q1010 of all three protomers. The backbone atoms (CA,C,N,O) of the rest of the system (Q14-R685,S686-L962,L1012-V1164) were restrained using 10.0 kcal/(mol Å$^2$) positional restraint. The RMSD was reduced from 9.4 Å to 0 using 100000 kcal/(mol Å$^2$) restraint during 30 ns of SMD at 310 K. After SMD, the system was relaxed during 30 ns MD at 310 K while applying 0.1 kcal/(mol Å$^2$) positional restraints to all CA atoms.

*Nudged elastic band optimization of the CH extension pathway in the 3-up spike*

We used the partial nudged elastic band (NEB) method(8) as implemented(9) on GPUs in Amber to generate a minimum energy path for the CH extension process. The 3-up spike structure from MD initiated from 7CAK and 3-up spike with extended central helices (see above) served as the two endpoints, which remain static throughout the calculation. Thirty additional beads were simulated in parallel between these two endpoints, with the first 16 initiated from the first endpoint and the last 16 from the second endpoint. The NEB springs were applied with a 1 kcal/ (mol * Å$^2$) force constant to the N, CA and C atoms of the entire protein, as well as the carbon atoms of the glycans attached to N234 since previous work(10) implicated this glycan site in RBD dynamics. NEB was run in a four-step protocol that involved heating, equilibrating, annealing, and MD. The set of 32 images were first heated from 100 to 300 K over 0.5 ns at constant volume, followed by 1 ns of equilibration at constant pressure of 1 bar and temperature of 300K. The system was then annealed by heating to 400 K over 2 ns, maintaining 400 K for 1 ns, cooling to 300 K over 2 ns, followed by a final 10 ns at 300 K to generate the final path. Following MD, a further 240ns of fully unrestrained MD was performed for the 3-up spike with extended CH.

*Energy postprocessing of simulation snapshots*

The solvation free energy of R1000 was estimated using Poisson-Boltzmann calculations, with a similar strategy as we reported(11) previously. Only the partial charges of the three R1000 were kept (including backbone and sidechains), while the partial charges of all other atoms were set to zero. The Amber *PBSA* module was used for PB calculations on the full spike. Default options were used except a grid size of 1 Å to reduce the memory requirements. Mbondi(12) atomic radii were used. Dielectric constants used in PB calculation were 78.5 and 1 for solvent and solute, respectively. Snapshots spaced 10 ns apart were taken from 250 ns of each simulation, resulting in 25 frames each for closed, 3-up and 3-up extended simulations, respectively.

*Calculation of the UH rotation angle*

We defined four Center of Mass (CoM) points to measure a dihedral that quantifies the UH rotation for each protomer. Amino acids N1054, S1055, G1060, and V1061 in the connector domain were used to calculate the first CoM point. The second point included T778, N779, E780, V781 (UH), the third point included R765 to T768 (UH) and the fourth point included T747 to S750 (UH). Only the Cα atoms were used to calculate the CoM locations. The dihedral values were calculated using cpptraj(13).

*Analysis of structures from the PDB*

81 structures were extracted from the PDB (**Table S1**). Only SARS-CoV-2 spike trimers including both the S1 and S2 subunits were included, except for the post-fusion 6XRA which includes only the S2 subunit. No model building was performed. If any atoms were missing from a structure, it was not included in that particular analysis. Chain information was corrected in some structures such that protomers were counterclockwise as chains A, B and C. PDB files with amino acid position numbering that did not correspond to the Wuhan sequence (e.g. 986/987 at the CH top) were excluded.

The RBD to HR1 distance was defined as the distance from the N atom of S383 to the O atom of R983. These amino acids have been replaced successfully with a disulfide bond(14-16), substantiating their close contact. The RBD to CH distance was defined as the distance between CA atoms of D427 and K986. Using the RBD-HR1 distance, **159 protomers exhibited a closed RBD, and 78 protomers exhibited an open RBD.** Average values are reported, with uncertainties reflecting the standard deviation of the distribution.

Salt bridge distances were calculated as the minimum value of six distances calculated between the two carboxyl oxygen atoms of Asp or Glu side chains and the three nitrogen atoms in the guanidino group of Arg. Asp to Ser distances were calculated as the minimum distance from either carboxyl oxygen to the hydroxyl oxygen of Ser. R1000 to S975 or I742 hydrogen bonding was calculated as the minimum distance of the three guanidino nitrogen atoms to the backbone oxygen of S975/I742. Uncertainties quoted on data from this structure set indicates the standard deviation over the measurement in all of the PDB structures.

*Water Density Analysis*

The grid command of cpptraj(13) was used to analyze the water density around R1000. 4200 frames from 420 ns of MD simulation of the closed model were taken for analysis. The structures were first aligned by overlapping residue V963-S1003 of the first protomer. Then the oxygen atoms of water molecules within a cubic region which centered at CE1 atom of F970 of the first protomer with edge length of 8 Å were counted using a bin width of 0.5 Å. The grid was visualized in VMD(17).

*Miscellaneous*

All structure images in this work were made using VMD(17) version 1.9.5a4. Structural analysis of the PDB structures and MD simulations was performed with cpptraj(13).

*References*


1. Y. Cai *et al.*, Distinct conformational states of SARS-CoV-2 spike protein. *Science* **369**, 1586-1592 (2020).
2. L. Fallon *et al.*, Free Energy Landscapes for RBD Opening in SARS-CoV-2 Spike Glycoprotein Simulations Suggest Key Interactions and a Potentially Druggable Allosteric Pocket. *ChemRxiv* **Preprint** (2020).
3. Y. Watanabe, J. D. Allen, D. Wrapp, J. S. McLellan, M. Crispin, Site-specific glycan analysis of the SARS-CoV-2 spike. *Science* **369**, 330-333 (2020).
4. B. Turoňová *et al.*, In situ structural analysis of SARS-CoV-2 spike reveals flexibility mediated by three hinges. *Science* **370**, 203-208 (2020).
5. T. Zhou *et al.*, Cryo-EM Structures of SARS-CoV-2 Spike without and with ACE2 Reveal a pH-Dependent Switch to Mediate Endosomal Positioning of Receptor-Binding Domains. *Cell Host & Microbe* **28**, 867-879.e865 (2020).
6. C. M. Carr, C. Chaudhry, P. S. Kim, Influenza hemagglutinin is spring-loaded by a metastable native conformation. *Proceedings of the National Academy of Sciences* **94**, 14306 (1997).
7. C. M. Carr, P. S. Kim, A spring-loaded mechanism for the conformational change of influenza hemagglutinin. *Cell* **73**, 823-832 (1993).
8. C. Bergonzo, A. J. Campbell, R. C. Walker, C. Simmerling, A partial nudged elastic band implementation for use with large or explicitly solvated systems. *International Journal of Quantum Chemistry* **109**, 3781-3790 (2009).
9. D. Ghoreishi, D. S. Cerutti, Z. Fallon, C. Simmerling, A. E. Roitberg, Fast Implementation of the Nudged Elastic Band Method in AMBER. *Journal of Chemical Theory and Computation* **15**, 4699-4707 (2019).
10. L. Casalino *et al.*, Beyond Shielding: The Roles of Glycans in the SARS-CoV-2 Spike Protein. *ACS Central Science* **6**, 1722-1734 (2020).
11. H. Nguyen, D. R. Roe, C. Simmerling, Improved Generalized Born Solvent Model Parameters for Protein Simulations. *Journal of Chemical Theory and Computation* **9**, 2020-2034 (2013).
12. A. Onufriev, D. Bashford, D. A. Case, Modification of the Generalized Born Model Suitable for Macromolecules. *The Journal of Physical Chemistry B* **104**, 3712-3720 (2000).

13. D. R. Roe, T. E. Cheatham, PTRAJ and CPPTRAJ: Software for Processing and Analysis of Molecular Dynamics Trajectory Data. *Journal of Chemical Theory and Computation* **9**, 3084-3095 (2013).
14. X. Xiong *et al.*, A thermostable, closed SARS-CoV-2 spike protein trimer. *Nature Structural & Molecular Biology* **27**, 934-941 (2020).
15. M. McCallum, A. C. Walls, J. E. Bowen, D. Corti, D. Veesler, Structure-guided covalent stabilization of coronavirus spike glycoprotein trimers in the closed conformation. *Nature Structural & Molecular Biology* **27**, 942-949 (2020).
16. R. Henderson *et al.*, Controlling the SARS-CoV-2 spike glycoprotein conformation. *Nature Structural & Molecular Biology* **27**, 925-933 (2020).
17. W. Humphrey, A. Dalke, K. Schulten, VMD: visual molecular dynamics. *J Mol Graph* **14**, 33-38, 27-38 (1996).
